# Mass mortality at penguin mega-colonies due to avian cholera confounds H5N1 HPAIV surveillance in Antarctica

**DOI:** 10.64898/2025.12.16.694678

**Authors:** Matteo Iervolino, Anne Günther, Leonard Schuele, Florencia Soto, Lonneke Leijten, Theo M. Bestebroer, Adam Coerper, Alice Reade, Ben Wallis, Simeon Lisovski, Martin Beer, Meagan Dewar, Thijs Kuiken, Lineke Begeman, Ralph E.T. Vanstreels

**Affiliations:** Department of Viroscience, Erasmus University Medical Center; Rotterdam, the Netherlands; Institute of Diagnostic Virology, Friedrich-Loeffler-Institut, Federal Research Institute for Animal Health; Greifswald-Insel Riems, Germany; Instituto de Biología de Organismos Marinos (IBIOMAR-CONICET); Puerto Madryn, Argentina; Ocean Expeditions Support Vessel S/V Australis; Sydney, New South Wales, Australia; Alfred Wegener Institute Helmholtz Centre for Polar and Marine Research; Potsdam, Germany; Future Regions Research Centre, Federation University Australia; Berwick, Victoria, Australia; Karen C. Drayer Wildlife Health Center, One Health Institute, School of Veterinary Medicine, University of California, Davis; Davis, CA, USA

## Abstract

In the austral summer 2023/2024, H5N1 high pathogenicity avian influenza virus (HPAIV) was reported for the first time in Antarctica. Concerns of HPAIV causing high mortality of seabirds and mammals prompted immediate efforts to track its spread and impact on endemic wildlife. In March 2024, we visited the Danger Islands archipelago, that hosts two mega-colonies of Adélie penguins, and observed an unusual mortality estimated in thousands of Adélie penguins and other species. Swabs and tissues were collected for molecular detection of infectious agents from 49 carcasses, and additional tissues for histology from a selection of 9 carcasses. We unexpectedly detected *Pasteurella multocida* DNA in 46 of 49 individuals, and diagnosed avian cholera, and not HPAI, as the cause of death of most of these animals. By metagenomics, we retrieved the genomic sequences of the *Pasteurella multocida* strain which caused the epizootic, and the phylogenetic analysis showed a close relation with strains previously reported in the Southern Ocean area. This study confirms avian cholera as a relevant cause of mortality in the Antarctic region, and overall highlights the importance of considering avian cholera in the differential diagnoses during mortality events in Antarctica, even with the concurrent circulation of HPAIV.

## Introduction

Infectious diseases are one of the main causes of mortality in wildlife worldwide. Despite being geographically relatively isolated from other animal communities and their pathogens, wildlife in Antarctica has been reported to die from infectious diseases [1–6]. However, limited effort has been spent in conducting comprehensive investigations regarding causative agents, host and geographic range and prevalence of Antarctic wildlife diseases. Although mass mortalities of wildlife have been sporadically reported in Antarctica, the remoteness and harsh environment have limited the detailed investigation of the causes of many such events [1,7–9]. Influenza A viruses (IAV) are known to circulate in Antarctic wildlife but, until recently, all strains detected were of low pathogenicity avian influenza (LPAI) and were not associated with significant morbidity or mortality [10–13]. In the summer of 2023/2024, high pathogenicity avian influenza virus (HPAIV) of the subtype H5N1 clade 2.3.4.4b was reported for the first time in the Antarctic Peninsula, causing significant wildlife mortality [5,6,14,15].

Penguins are flagship species of Antarctica, and there has been great international concern on whether these species, especially those already challenged by climate and environmental changes, could be impacted by the arrival of H5 HPAIV to the region, also considering that this virus has caused significant mortality of wild and captive penguins elsewhere [4,16–20]. Although H5 HPAIV RNA was detected in the brain of Adélie (*Pygoscelis adeliae*) and gentoo penguins (*Pygoscelis papua*) found dead throughout the Antarctic Peninsula in 2023/2024, detailed pathological investigation did not corroborate HPAIV infection as the cause of death, suggesting that those penguins had died from other causes [6]. RNA from H5 viruses was also detected in oropharyngeal and cloacal swabs from apparently healthy Adélie penguins at various sites in the Antarctic Peninsula in 2023/2024 [21]. However, it was not determined whether the strains were LPAIV or HPAIV [22]. Therefore, to which extent H5N1 HPAIV has circulated and represented a significant cause of mortality of Antarctic penguins remains unclear.

Four penguin species have substantial breeding populations in the Antarctic Peninsula: Adélie (1,204,000 nests), chinstrap (*Pygoscelis antarcticus*; 650,000 nests), gentoo (173,000 nests), and macaroni penguins (*Eudyptes chrysolophus*; 2,000 nests) [23,24]. Adélie penguins, the most abundant of the Antarctic penguins, have experienced substantial changes in their population size and distribution over recent decades [25–31]. However, despite the overall decrease in the Antarctic Peninsula region, the Adélie penguin population at the Danger Islands, an archipelago of seven small islands in the northern Weddell Sea (Figure 1), appears to have remained stable at least until 2015, representing an important sanctuary for the species in the region [32]. Although relatively small, the archipelago has an outstandingly high abundance and diversity of seabirds, including two mega-colonies of Adélie penguins, which has recently prompted its designation as an Antarctic Specially Protected Area (ASPA no. 180) [33]. Combined, the Danger Islands represent 55% of the species’ population in the Antarctic Peninsula [24,32]. In addition to Adélie penguins, the islands are home to gentoo penguins and chinstrap penguins, as well as shags, petrels, skuas, gulls, and sheathbills [32,33].

**Figure 1.**
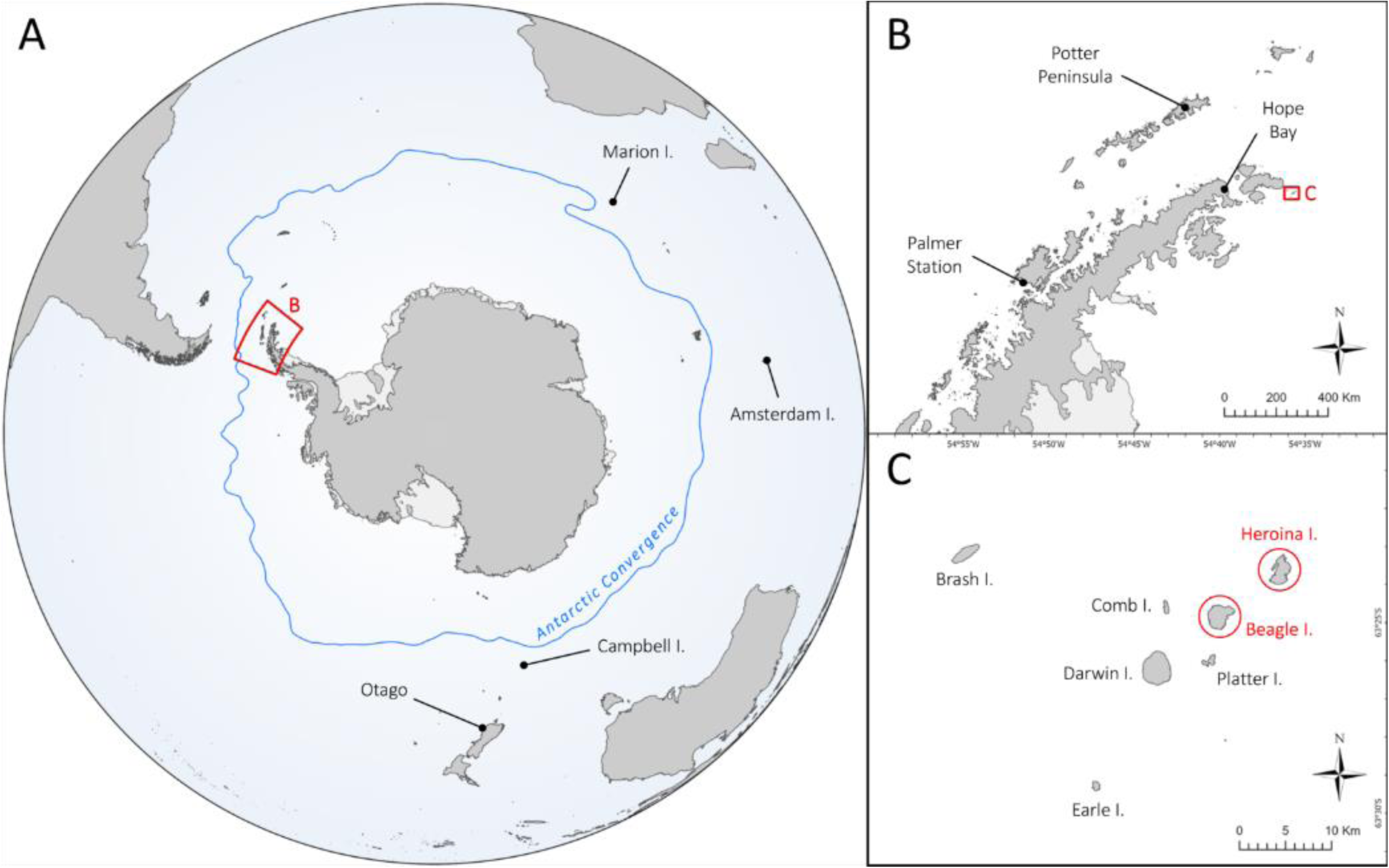
Location of the study sites (Heroina and Beagle Islands) and other sites mentioned in the text. **A)** Southern hemisphere with Antarctic Convergence indicated in blue. **B)** Antarctic Peninsula. **C)** Danger Islands, with Heroina and Beagle Island (part of this study) highlighted in red.

Mass mortality events of Adélie penguins have been reported in Antarctica sporadically over the last century, with the majority of incidents related to environmental factors [25,34,35]. However, in the austral summers of 1999/2000 and 2000/2001, avian cholera was diagnosed as the cause of mass mortalities of Adélie penguins and other seabirds at Hope Bay, Antarctic Peninsula [3]. Additionally, avian cholera was repeatedly reported to cause mortality in seabirds in the Antarctic and Subantarctic regions between 1979 and 2024 [2,3,6,36–41]. Avian cholera –also known as pasteurellosis or fowl cholera– is an infectious disease caused by the Gram-negative bacterium *Pasteurella multocida,* which is classified based on five capsular groups (A, B, D, E, F) and eight lipopolysaccharide (LPS) group loci (L1-L8), which are responsible for the expression of 16 LPS types (1-16) [42]. Avian cholera is a common cause of acute septicemic and respiratory disease in birds worldwide, with natural infection detected in more than 190 bird species [43]. Whereas it frequently occurs in domestic birds in most countries, reports of avian cholera in wild birds have been limited, except for wild waterfowl in North America, where large epizootics have been annually reported since the 1940s [43–46]. As most wild birds die acutely from avian cholera, they are usually found already dead and in good nutritional condition, but clinical signs may involve lethargy, nasal discharge, and neurological signs. Post-mortem findings usually include hyperemia, hemorrhages, and necrotic foci, and, consistent with septicemia, massive numbers of small coccoid bacteria are visible in blood vessels and tissues [43,45].

In March 2024, during the HPAI Australis Expedition, set up to study the spread of H5 HPAIV and its impact on wildlife in the Antarctic Peninsula [14], we visited Heroina and Beagle Island (Danger Islands). This decision was based on previous findings from December 2023, when eight dead brown skuas were found on the main landing ramp of Heroina Island, with oral swabs testing negative for IAV at the APHA reference laboratory in the United Kingdom [47,48]. Also in December 2023, cloacal swabs were collected from 16 healthy adult Adélie penguins at Beagle Island, seven of which were positive for H5 viruses [21,22]. At the time of our visit, we found hundreds of Adélie penguins deceased on both islands. To investigate their cause of death, we collected swabs and tissues from dead animals, and performed autopsies. Here, we present the results of our virological, bacteriological and histopathological investigations, and the surprising finding of avian cholera as the probable cause of death of these mortalities. We also show a detailed characterization of the *P. multocida* strains found on Beagle and Heroina Island as well as those found at other sites visited in the same context [6], and phylogenetic analyses we employed to study the possible incursion and transmission pathways. These results ultimately support the need to consider alternative diagnoses such as avian cholera, even with concurrent detection of HPAIV, and provide methodological tools for this purpose.

## Results

### Field observations

During our survey at Heroina Island, a total of 537 Adélie penguin carcasses (172 adults and 365 chicks, most of which were near fledging) were counted at the island’s main landing ramp (red area in Figure 2A). Additionally, it was estimated that several hundreds, possibly thousands, of Adélie penguin carcasses were also present in the island’s north plateau (orange areas in Figure 2A). Furthermore, the carcasses of four adult brown skuas (*Stercorarius antarcticus*), seven unidentified skuas (*Stercorarius* sp.; 1 juvenile, 6 adults), three snowy sheathbills (*Chionis albus*; undetermined age class), one southern giant petrel (*Macronectes giganteus*; undetermined age class), and one Antarctic fur seal (*Arctocephalus gazella*; undetermined age class) were found (Figure 2A). These likely included the skuas that had been found earlier in December 2023. All carcasses were frozen and partially covered by snow, and in moderate to advanced state of autolysis (Table 1), suggesting mortality may have occurred several weeks prior to our visit. Relatively small numbers of live animals were present at the time of the visit: roughly 40 gentoo penguins, 40 Antarctic fur seals, 20 Adélie penguins, 10 Antarctic shags (*Leucocarbo bransfieldensis*), four Weddell seals (*Leptonychotes weddellii*), and an undefined number of kelp gulls (*Larus dominicanus*). The paucity of live Adélie penguins was not unexpected given the timing of the visit; Adélie penguins breed from October to February [23]. All live animals were apparently healthy, except for one juvenile Antarctic fur seal showing signs of lethargy, conjunctivitis, and respiratory distress (video in Supplementary File 1).

**Figure 2.**
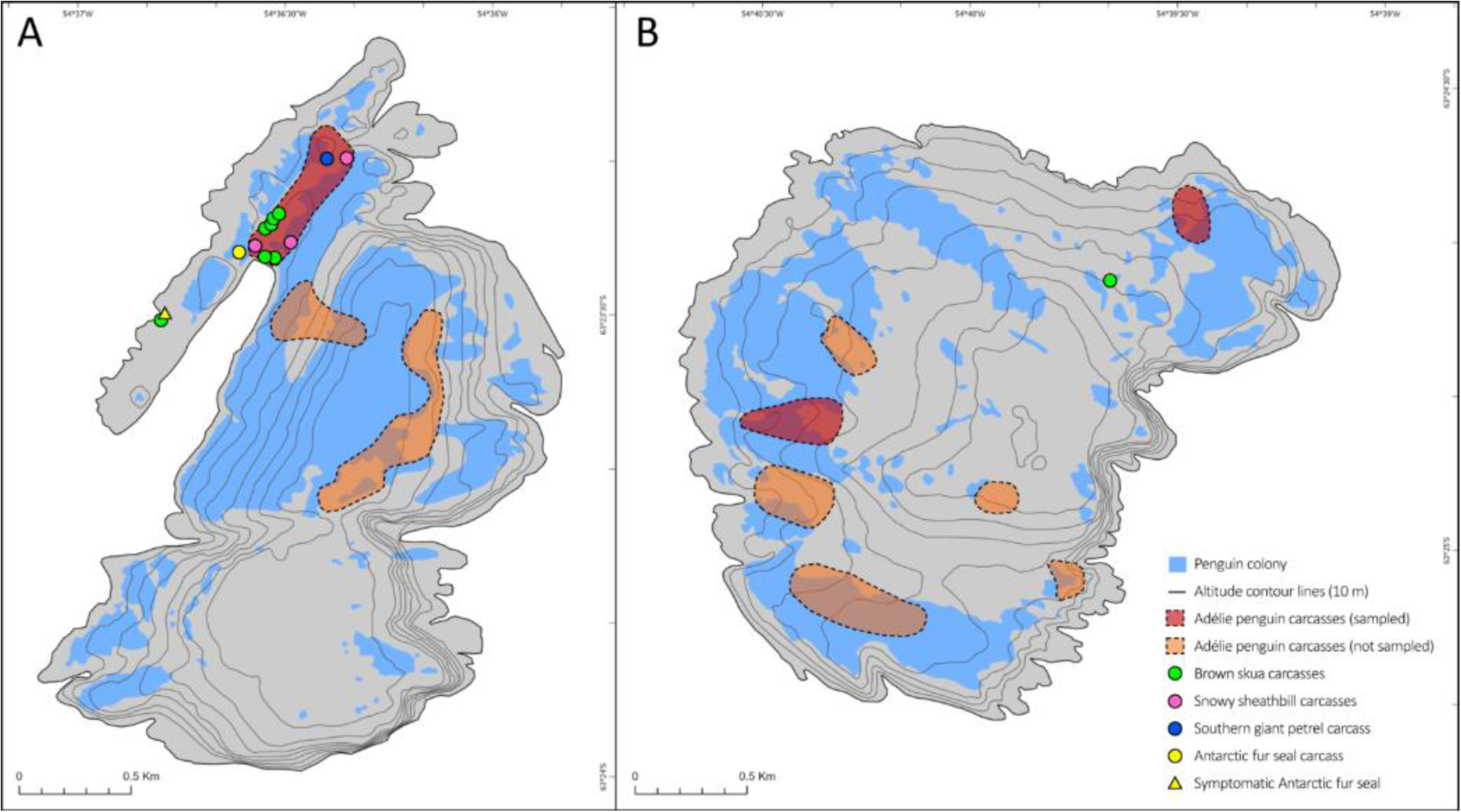
Spatial distribution of wildlife mortality observed at the Danger Islands, March 2024. **A)** Heroina Island. **B)** Beagle Island. Island outlines, penguin colonies and altitude contours were re-drawn from ATCM (2024).

**Table 1.** Summary of mortality and overview of diagnostic criteria for influenza A virus (IAV) infection and avian cholera (*Pasteurella multocida*, Pm), and diagnoses of animals found dead. The included diagnostic criteria are the influenza A virus RNA loads (IAV RNA) from the *M* gene RT-qPCR, the influenza A virus NP antigen immunohistochemistry (IAV NP IHC), the *P. multocida* DNA loads from the *kmt1* qPCR (Pm DNA), and the histology results (Pm histo; H&E and ISH). For the species category “Skua”, the ID is indicated as BSK (Brown skua, *Stercorarius antarcticus*) if the confidence level for species identification was ≥75%. The confidence level of the diagnosis applies to all diagnoses within the same animal.

| Location | Species | ID | Age class | Decomposition and scavenging state | Nutritional condition | IAV |  | Avian cholera |  | Diagnosis |  |
| --- | --- | --- | --- | --- | --- | --- | --- | --- | --- | --- | --- |
|  |  |  |  |  |  | IAV RNA (tissue) | IAV NP IHC | Pm DNA, brain | Pm histo | Diagnosis at death in order of severity | Confidence level |
| Heroína Island | Adélie penguin (*Pygoscelis adeliae*) | AP27 | Ad | Moderate | Good | - | - # | +++ | + | 1. Avian cholera | High |
|  |  | AP28 | Ad | Moderate | Poor | - | na | +++ | na | 1. Avian cholera | Medium |
|  |  | AP29 | Ad | Moderate | Good | - | - | +++ | + | 1. Avian cholera | High |
|  |  | AP30 | Ad | Advanced | Good | - | na | +++ | na | 1. Avian cholera | Medium |
|  |  | AP31 | Ad | Advanced | Good ‡ | + (L) | na | +++ | na | 1. Avian cholera; 2. Avian Influenza | Medium |
|  |  | AP32 | Ad | nd | Good ‡ | - | na | +++ | na | 1. Avian cholera | Medium |
|  |  | AP33 | Chick | Moderate | Poor | - | - | ++ | + | 1. Avian cholera; 2. Starvation | Medium |
|  |  | AP34 | Chick | Advanced | nd | - | na | ++ | na | 1. Avian cholera | Medium |
|  |  | AP35 | Chick | Moderate | Poor | - | na | + | na | 1. Starvation; 2. Avian cholera | Medium |
|  |  | AP36 | Chick | Moderate | nd | - | na | + | na | No diagnosis | Inconclusive |
|  |  | AP37 | Ad | Moderate | Good | + (O, I, LI, SF) | - | + | - | No diagnosis | Inconclusive |
|  |  | AP38 | Ad | Moderate, Partly scavenged | Good | + (I, LI, SF) | - | +++ | + | 1. Avian cholera; 2. Avian influenza | High |
|  |  | AP39 | Ad | Moderate | nd | - | na | +++ | na | 1. Avian cholera | Medium |
|  |  | AP40 | Ad | Moderate | nd | - | na | +++ | na | 1. Avian cholera | Medium |
|  |  | AP41 | Ad | Moderate | nd | + (L) | na | +++ | na | 1. Avian cholera; 2. Avian influenza | Medium |
|  |  | AP42 | Chick | Moderate | nd | - | na | + | na | No diagnosis | Inconclusive |
|  |  | AP43 | Chick | Moderate | nd | - | na | - | na | No diagnosis | Inconclusive |
|  |  | AP44 | Chick | Moderate | nd | - | na | - | na | No diagnosis | Inconclusive |
|  |  | AP45 | Chick | Moderate | nd | - | na | - | na | No diagnosis | Inconclusive |
|  |  | AP46 | Chick | Moderate | nd | - | na | - | na | No diagnosis | Inconclusive |
|  |  | AP47 | Chick | Moderate | nd | + (B, L) | na | - | na | No diagnosis | Inconclusive |
|  | Skua (*Stercorarius sp.*) | SK18 | Ad | Advanced | nd | - | na | +++ | na | 1. Avian cholera | Medium |
|  |  | SK19 | Ad | Advanced | nd | + (B) | na | ++ | na | 1. Avian cholera; 2. Avian Influenza | Medium |
|  |  | BSK20 | Ad | Advanced | nd | - | na | +++ | na | 1. Avian cholera | Medium |
|  |  | BSK21 | Ad | Advanced | nd | - | na | ++ | na | 1. Avian cholera | Medium |
|  |  | SK22 | Juv | Advanced | nd | - | na | +++ | na | 1. Avian cholera | Medium |
|  |  | SK23 | Ad | Advanced | nd | + (B) | na | ++ | na | 1. Avian cholera; 2. Avian influenza | Medium |
|  |  | SK24 | Juv | Advanced | nd | + (L) | na | + | na | 1. Avian cholera; 1. Avian influenza | Low |
|  |  | BSK25 | Ad | Advanced | nd | - | na | ++ | na | 1. Avian cholera | Medium |
|  |  | SK26 | Ad | Advanced | nd | - | na | ++ | na | 1. Avian cholera | Medium |
|  |  | SK27 | Juv | Advanced | nd | - | na | ++ | na | 1. Avian cholera | Medium |
|  | Snowy sheathbill (*Chionis albus*) | SB02 | nd | Advanced | nd | - | - | + | - | No diagnosis | Inconclusive |
|  |  | SB03 | nd | Advanced | nd | - | na | +++ | na | 1. Avian cholera | Medium |
|  |  | SB04 | nd | Advanced | nd | - | na | ++ | na | 1. Avian cholera | Medium |
|  | Southern giant petrel (*Macronectes giganteus*) | GP01 | Ad | Advanced | nd | - | na | + | na | 1. Avian cholera | Low |
|  | Antarctic fur seal (*Arctocephalus gazella*) | FS03* | Juv | na | Poor | -(swab)* | na | na | na | No diagnosis | Inconclusive |
|  |  | FS04 | nd | Advanced | nd | - | na | ++ | na | No diagnosis | Inconclusive |
| Beagle Island | Adélie penguin (*Pygoscelis adeliae*) | AP48 | Ad | Moderate | Good | - | - | +++ | + | 1. Avian cholera | High |
|  |  | AP49 | Chick | Scanvenged | nd | - | na | +++ | na | 1. Avian cholera | Medium |
|  |  | AP50 | Ad | Scanvenged | Good † | - | - | +++ | + | 1. Avian cholera | High |
|  |  | AP51 | Ad | Moderate | Moderate | - | - | +++ | + | 1. Avian cholera | Medium |
|  |  | AP52 | Ad | Moderate | Good | - | na | +++ | na | 1. Avian cholera | Medium |
|  |  | AP53 | Ad | Moderate | Good | - | na | +++ | na | 1. Avian cholera | Medium |
|  |  | AP54 | Ad | Moderate | Good | - | na | ++ | na | 1. Avian cholera | Medium |
|  |  | AP55 | Ad | Moderate | Good ‡ | - | na | +++ | na | 1. Avian cholera | Medium |
|  |  | AP56 | Ad | Moderate | Good ‡ | - | na | +++ | na | 1. Avian cholera | Medium |
|  |  | AP57 | Ad | Moderate | nd | - | na | +++ | na | 1. Avian cholera | Medium |
|  |  | AP58 | Ad | Moderate | Good ‡ | - | na | +++ | na | 1. Avian cholera | Medium |
|  |  | AP59 | Ad | Moderate | nd | - | na | +++ | na | 1. Avian cholera | Medium |
|  | Skua (*Stercorarius sp.*) | SK28 | Ad | Moderate | Good | - | na | ++ | na | 1. Avian cholera | Medium |

IAV RNA and Pm DNA: *M* RT-qPCR and *kmt1* (RT-)qPCR DNA loads (40 - Cq): -, not detected; +, 1 to 8; ++, 9 to 16; +++, 17 to 40. Tissues: B, brain; L, lung; O, oropharyngeal swab; I, intestine; LI, liver; SF, skin or feather follicle.
\* FS03 was alive. The IAV RT-qPCRs and kmt1 qPCR were performed on nasal discharge and per-anal swabs.

At Beagle Island, hundreds of Adélie penguin carcasses (approximately 40–50% adults and 50–60% chicks, most of which were near fledging) were observed, distributed on various sub-colonies of the island (red and orange areas in Figure 2B). Additionally, an adult unidentified skua carcass was found on the northeast of the island (Figure 2B). Heavy snowfall and limited time on land due to navigation safety concerns constrained our ability to further quantify mortality and live animals at this island.

Overall, adult Adélie penguin carcasses at both islands appeared to be in a good nutritional state, showing ample fat stores and well-developed muscles, and limited scavenging. A subset of 33 dead Adélie penguins was selected for sample collection (11 adults and 10 chicks from Heroina Island and 11 adults and 1 chick from Beagle Island). Of these, five Adélie penguins (AP27, AP29, AP33, AP37, AP38) from Heroina Island and three Adélie penguins (AP48, AP50, AP51) from Beagle Island were fully autopsied. All carcasses of other species mentioned above were also sampled.

### Bacteriological and virological detections

#### Low influenza A virus RNA loads in swabs and tissues from carcasses

Low IAV RNA loads were detected by RT-qPCR (*M* gene) in samples from 5/21 Adélie penguins and 3/10 skuas from Heroina Island (Table 1). In Adélie penguins, the viral RNA loads (40 – Cq) ranged between 3.4 and 7.9, while in skuas between 3.6 and 6.4. None of the carcasses from other species (fur seals, sheathbills, and giant petrel) on Heroina Island and no carcasses from Beagle Island tested positive for IAV (Table 2 and Supplementary Table 1). Two (AP37 and AP47) of the IAV-positive penguins from Heroina Island tested positive in the H5 RT-qPCR, of which one (AP37) was also positive in the N1 RT-qPCR. None of the IAV- or H5-positive samples tested positive in the HPAIV RT-qPCR (Supplementary Table 1).

**Table 2.** Influenza A virus (IAV) loads (40-Cq, *M* gene) and *P. multocida* (Pm) bacterial loads (40-Cq, *kmt1* gene) in swabs and tissues of animals found dead from Heroina and Beagle Island. The IDs from onboard testing are indicated to compare with previous results [14,49].

| Location | Species | ID | ID onboard test [49] | Oropharyngeal swab |  | Cloacal swab |  | Brain |  | Lung |  | Intestine |  | Liver |  | Spleen |  | Kidney |  | Feather follicle |  |
| --- | --- | --- | --- | --- | --- | --- | --- | --- | --- | --- | --- | --- | --- | --- | --- | --- | --- | --- | --- | --- | --- |
|  |  |  |  | IAV | Pm | IAV | Pm | IAV | Pm | IAV | Pm | IAV | Pm | IAV | Pm | IAV | Pm | IAV | Pm | IAV | Pm |
| Heroina Island | Adélie penguin (*Pygoscelis adeliae*) | AP27 | ADPE_1_Hero | na | na | na | na | nd | 17.2 | nd | 18.7 | na | na | na | na | na | na | na | na | na | na |
|  |  | AP28 | ADPE_2_Hero | na | na | na | na | nd | 20.3 | nd | 19.9 | na | na | na | na | na | na | na | na | na | na |
|  |  | AP29 | ADPE_3_Hero | na | na | na | na | nd | 19.1 | nd | 20.2 | na | na | na | na | na | na | na | na | na | na |
|  |  | AP30 |  | na | na | na | na | nd | 19.2 | nd | 20.2 | na | na | na | na | na | na | na | na | na | na |
|  |  | AP31 |  | na | na | na | na | nd | 17.4 | 7.5 | 18.6 | na | na | na | na | na | na | na | na | na | na |
|  |  | AP32 |  | na | na | na | na | nd | 19.0 | nd | 21.0 | na | na | na | na | na | na | na | na | na | na |
|  |  | AP33 |  | na | na | na | na | nd | 10.7 | nd | 13.0 | na | na | na | na | na | na | na | na | na | na |
|  |  | AP34 |  | na | na | na | na | nd | 10.9 | nd | 10.3 | na | na | na | na | na | na | na | na | na | na |
|  |  | AP35 | ADPE_4_Hero | na | na | na | na | nd | 8.4 | nd | 10.0 | na | na | na | na | na | na | na | na | na | na |
|  |  | AP36 |  | na | na | na | na | nd | 7.0 | nd | 10.0 | na | na | na | na | na | na | na | na | na | na |
|  |  | AP37 | ADPE_8_Hero | 6.4 | 8.6 | nd | 10.2 | nd | 5.5 | nd | 8.6 | 4.7 | 10.5 | 7.9 | 11.8 | nd | 3.8 | nd | 3.9 | 7.3 | 10.7 |
|  |  | AP38 |  | nd | 14.1 | na | na | nd | 20.6 | nd | 21.3 | 4.9 | 23.4 | 5.3 | 22.6 | na | na | nd | 20.4 | 5.2 | 14.3 |
|  |  | AP39 | ADPE_9_Hero | na | na | na | na | nd | 20.3 | nd | 21.5 | na | na | na | na | na | na | na | na | na | na |
|  |  | AP40 | ADPE_10_Hero | na | na | na | na | nd | 19.1 | nd | 20.0 | na | na | na | na | na | na | na | na | na | na |
|  |  | AP41 | ADPE_11_Hero | na | na | na | na | nd | 19.7 | 3.4 | 19.5 | na | na | na | na | na | na | na | na | na | na |
|  |  | AP42 |  | na | na | na | na | nd | 4.5 | nd | 3.9 | na | na | na | na | na | na | na | na | na | na |
|  |  | AP43 | ADPE_5_Hero | na | na | na | na | nd | nd | nd | 4.1 | na | na | na | na | na | na | na | na | na | na |
|  |  | AP44 |  | na | na | na | na | nd | nd | nd | nd | na | na | na | na | na | na | na | na | na | na |
|  |  | AP45 |  | na | na | na | na | nd | nd | nd | nd | na | na | na | na | na | na | na | na | na | na |
|  |  | AP46 | ADPE_6_Hero | na | na | na | na | nd | nd | nd | nd | na | na | na | na | na | na | na | na | na | na |
|  |  | AP47 | ADPE_7_Hero | na | na | na | na | 7.7 | nd | 5.7 | 3.6 | na | na | na | na | na | na | na | na | na | na |
|  | Skua (*Stercorarius* sp.) | SK18 | Skua_1_Hero | na | na | na | na | nd | 19 | nd | 22.3 | na | na | na | na | na | na | na | na | na | na |
|  |  | SK19 | Skua_2_Hero | na | na | na | na | 3.6 | 15 | nd | 20.6 | na | na | na | na | na | na | na | na | na | na |
|  |  | BSK20 | Skua_3_Hero | na | na | na | na | nd | 16.7 | nd | 22.0 | na | na | na | na | na | na | na | na | na | na |
|  |  | BSK21 |  | na | na | na | na | nd | 15.2 | na | na | na | na | na | na | na | na | na | na | na | na |
|  |  | SK22 |  | na | na | na | na | nd | 17.3 | na | na | na | na | na | na | na | na | na | na | na | na |
|  |  | SK23 |  | na | na | na | na | 5.5 | 12.9 | na | na | na | na | na | na | na | na | na | na | na | na |
|  |  | SK24 | Skua_4_Hero | na | na | na | na | nd | 7.1 | 6.4 | 17.9 | na | na | na | na | na | na | na | na | na | na |
|  |  | BSK25 |  | na | na | na | na | nd | 12.7 | na | na | na | na | na | na | na | na | na | na | na | na |
|  |  | SK26 |  | na | na | na | na | nd | 11.6 | na | na | na | na | na | na | na | na | na | na | na | na |
|  |  | SK27 |  | na | na | na | na | nd | 10.8 | na | na | na | na | na | na | na | na | na | na | na | na |
|  | Snowy sheathbill (*Chionis albus*) | SB02 | SSB_3_Hero | na | na | na | na | nd | 4.4 | nd | 11.9 | na | na | na | na | na | na | na | na | na | na |
|  |  | SB03 | SSB_1_Hero | na | na | na | na | nd | 18.9 | na | na | na | na | na | na | na | na | na | na | na | na |
|  |  | SB04 | SSB_2_Hero | na | na | na | na | nd | 11.1 | na | na | na | na | na | na | na | na | na | na | na | na |
|  | Southern giant petrel (*Macronectes giganteus*) | GP01 | GP_1_Hero | na | na | na | na | nd | 6.6 | na | na | na | na | na | na | na | na | na | na | na | na |
|  | Antarctic fur seal (*Arctocephalus gazella*) | FS03.1* | AFS_alive_2_Hero | nd# | 5.9 | nd# | nd | na | na | na | na | na | na | na | na | na | na | na | na | na | na |
|  |  | FS03.2* | AFS_alive_2_Hero | nd# | nd | nd# | 3.2 | na | na | na | na | na | na | na | na | na | na | na | na | na | na |
|  |  | FS04 | AFS_1_Hero | na | na | na | na | nd | 14.2 | na | na | na | na | na | na | na | na | na | na | na | na |
| Beagle Island | Adélie penguin (*Pygoscelis adeliae*) | AP48 | ADPE_1_Beagle | nd | 15.3 | nd | 13.0 | nd | 21.8 | nd | 19.6 | nd | 17.0 | nd | 23.8 | na | na | nd | 21.8 | nd | 16.6 |
|  |  | AP49 | ADPE_2_Beagle | na | na | na | na | nd | 19.7 | nd | 18.6 | na | na | na | na | na | na | na | na | na | na |
|  |  | AP50 | ADPE_3_Beagle | na | na | na | na | nd | 18.2 | nd | 21.9 | na | na | na | na | na | na | na | na | na | na |
|  |  | AP51 | ADPE_4_Beagle | na | na | na | na | nd | 20.6 | nd | 22.3 | na | na | na | na | na | na | na | na | na | na |
|  |  | AP52 | ADPE_5_Beagle | na | na | na | na | nd | 21.3 | nd | 21.3 | na | na | na | na | na | na | na | na | na | na |
|  |  | AP53 | ADPE_6_Beagle | na | na | na | na | nd | 20.1 | nd | 21.0 | na | na | na | na | na | na | na | na | na | na |
|  |  | AP54 | ADPE_7_Beagle | na | na | na | na | nd | 12.6 | nd | 13.5 | na | na | na | na | na | na | na | na | na | na |
|  |  | AP55 | ADPE_8_Beagle | na | na | na | na | nd | 19.7 | na | na | na | na | na | na | na | na | na | na | na | na |
|  |  | AP56 | ADPE_9_Beagle | na | na | na | na | nd | 20.7 | na | na | na | na | na | na | na | na | na | na | na | na |
|  |  | AP57 | ADPE_10_Beagle | na | na | na | na | nd | 21.5 | na | na | na | na | na | na | na | na | na | na | na | na |
|  |  | AP58 | ADPE_11_Beagle | na | na | na | na | nd | 19.4 | nd | 21.8 | na | na | na | na | na | na | na | na | na | na |
|  |  | AP59 | ADPE_12_Beagle | na | na | na | na | nd | 20.8 | na | na | na | na | na | na | na | na | na | na | na | na |
|  | Skua (*Stercorarius* sp.) | SK28 | Skua_1_Beagle | na | na | na | na | nd | 12.5 | nd | 15.9 | na | na | na | na | na | na | na | na | na | na |

\* FS03 was alive and environmental oro-nasal and peri-anal secretions were sampled in two consecutive days.

To rule out possible inefficient viral RNA detection due to impaired RNA preservation in autolytic carcasses, all samples were tested for the presence of RNA from housekeeping genes (*GAPDH* for seabirds and *β-actin* for seals). Moderate to high *GAPDH* RNA loads (40 – Cq) were detected in seabird samples, ranging between 12.8 and 24.0 (mean 20.4 ± SD 1.7; n=101). Moderate *β-actin* RNA loads (16.4) were found in the brain sample of the deceased fur seal (Supplementary Table 2).

#### High *P. multocida* DNA loads in swabs and tissues from carcasses

*P. multocida* DNA was detected by qPCR (*kmt1* gene) of swabs and tissues from 30/33 Adélie penguins, 11/11 skuas, 3/3 snowy sheathbills, 1/1 Antarctic fur seal, and 1/1 southern giant petrel we sampled from Heroina and Beagle Islands. For each of the three Adélie penguins for which a full set of tissues was collected for molecular analyses (AP37, AP38, AP48), all samples tested positive for *P. multocida* DNA, indicating systemic infections. On average, the highest *P. multocida* DNA load (40 – Cq) was found in the liver (19.4 ± 6.6, n=3), followed by brain (17.0 ± 5.3, n=28), intestine (17.0 ± 6.4, n=3), and lungs (16.3 ± 6.2, n=26). In skuas, for which the liver was not sampled for any of the 11 individuals, the highest *P. multocida* DNA load was on average found in the lung (19.7 ± 2.8, n=5). A low *P. multocida* DNA load (13.6) was detected in the brain of a dead Antarctic fur seal from Heroina Island (Table 2, Supplementary Table 1).

#### Low *P. multocida* DNA loads in swabs from a live symptomatic fur seal

Environmental oronasal and peri-anal secretion swabs from one lethargic Antarctic fur seal at Heroina Island (FS03) tested positive for *P. multocida* DNA, with low bacterial DNA loads (5.9 and 3.2, respectively). No IAV RNA and low *β-actin* RNA loads were detected in the same swabs from this individual (Supplementary Table 1 and 2).

#### No *P. multocida* DNA loads in fecal samples from live animals

All 71 environmental fecal swabs collected from live and apparently healthy individuals (38 gentoo penguins, 14 kelp gulls, 13 Adélie penguins, 3 snowy sheathbills, 2 Antarctic fur seals, and 1 Weddell seal) from Heroina Island tested negative for *P. multocida* DNA and IAV RNA by (RT-)qPCR.

### Macroscopic and microscopic analyses

#### Causes of death were not obvious from macroscopic evidence

Autopsied carcasses were completely or partly frozen and were in moderate (sunken eyes, and all tissues except lungs having the same dark red color) to advanced (mummified remains) decomposition state (Table 1). Most penguins were in good nutritional condition. Lungs of four Adélie penguins were diffusely wet, and heavy, fitting with hyperemia and/or lung edema. No other macroscopic abnormalities were detected. One of the eight autopsied Adélie penguins had its stomach filled with krill (AP30), suggesting it had died peracutely.

#### Microscopic analysis corroborates systemic infection by *P. multocida* in Adélie penguins, but not in sheathbill

Organs from nine carcasses (5 Adélie penguins and 1 snowy sheathbill from Heroina Island, and 3 Adélie penguins from Beagle Island) were sampled for microscopic examination as tissues were deemed well enough preserved (Supplementary Table 3). For all tissues, freezing artefacts and autolysis hampered observation of more subtle lesions. Lesions typical for avian cholera were present in the livers of 5/6 Adélie penguin carcasses for which liver samples were collected (Supplementary Table 3). Whereas the hepatocytes generally showed poor cytoplasmic staining, loss of nuclei and vague cell borders due to autolysis, cells in the liver lesions were instead better preserved, and were characterized by randomly distributed areas of approximately 30–100 hepatocytes that had distinct cell borders, hypereosinophilic cytoplasm and showed loss of nuclei (coagulative necrosis). These areas contained a few heterophils and aggregates of small (1–2 μm diameter) coccoid bacteria, morphologically consistent with *P. multocida* (micro-abscesses) (Figure 3). *In situ* hybridization (ISH) targeting *P. multocida* DNA was used on the best-preserved carcasses of Adélie penguins from both islands (AP38 from Heroina Island and AP50 from Beagle Island). ISH confirmed the presence of *P. multocida* DNA co-localized with the lesions in both livers tested, as well as abundant *P. multocida* bacteria presence within liver sinusoids and blood vessels outside the lesions (Figure 3, Supplementary Figure 1).

**Figure 3.**
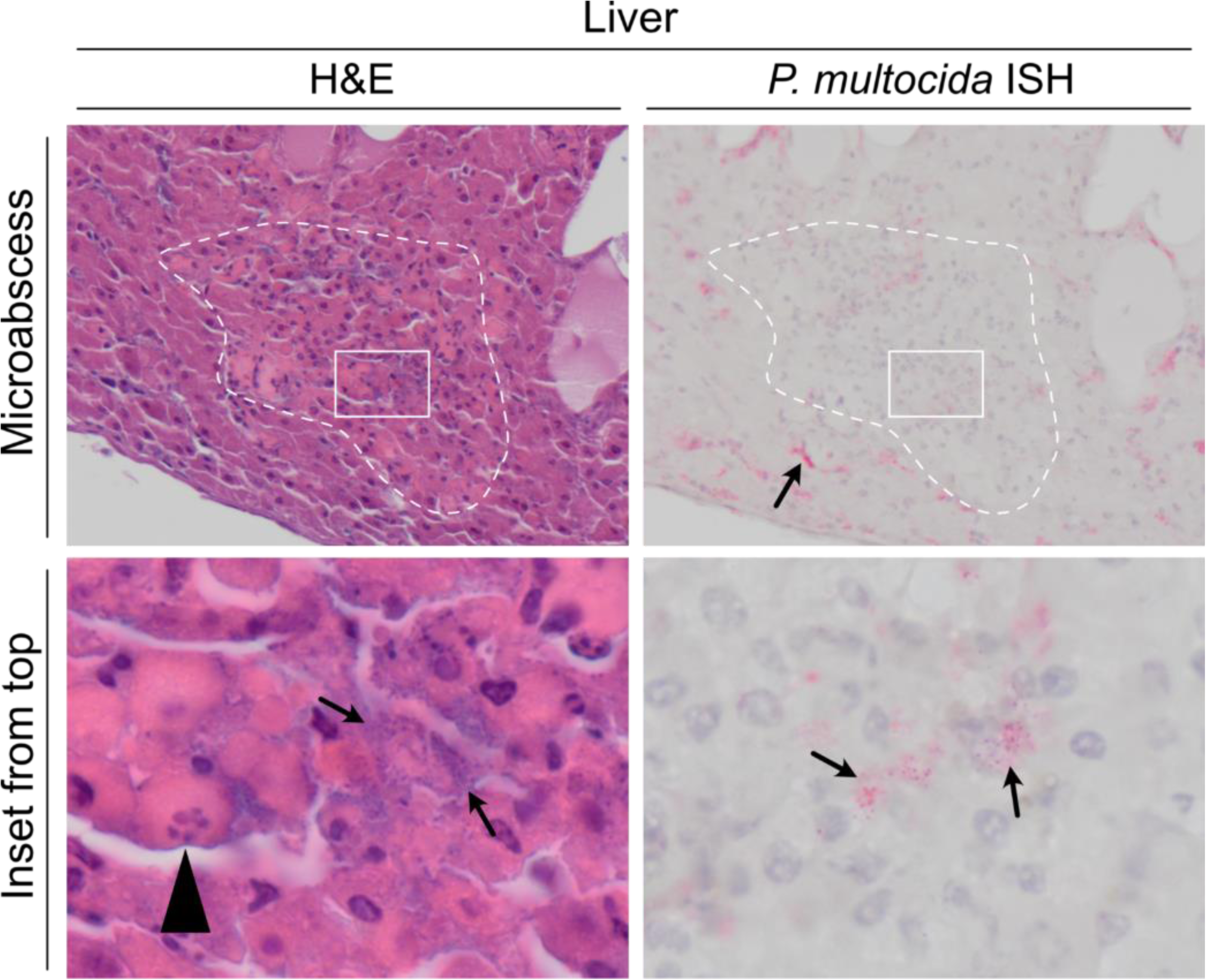
Serial sections of hematoxylin and eosin staining (H&E) and *P. multocida in situ* hybridization (ISH) of the liver of Adélie penguin. Liver of Adélie penguin (AP38) found dead. The white dotted line encircles the area with a microabscess, characterized by increased cellularity due to influx of inflammatory cells and hepatocellular necrosis (characterized by hypereosinophilic cytoplasm, karyorrhexis, pyknosis and karyolysis). The arrow points at the ISH-positive (staining pink) aggregates of bacteria within hepatic sinusoids. The inset below shows higher magnification of area within microabscess showing numerous small (1–2 µm diameter) coccoid bacteria morphologically consistent with *P. multocida*. The serial section in this area stains ISH-positive bacteria, confirming the presence of *P. multocida* in the lesion. The arrows point at *P. multocida*. The arrowhead points to a necrotic cell showing swelling, hypereosinophilia and karyorrhexis.

Besides these avian cholera-associated lesions in the liver, aggregations of bacteria morphologically consistent with *P. multocida*, and confirmed by ISH from AP38 and AP50, were variably observed in middle to large blood vessel of lungs, brain, trachea, heart and kidney (Figure 4, Supplementary Table 3). Occasionally these bacteria were admixed with fibrillar eosinophilic material, heterophils and macrophages attached to blood vessel walls (bacterial thrombi) (Figure 4). For two Adélie penguins (AP29 and AP51), no liver samples were available for microscopic analyses and qPCR. However, *Pasteurella-*like bacterial colonies were present in the blood vessels of lungs, with matching ISH staining for *P. multocida* (AP29). Overall, these results indicate multi-organ disease from *P. multocida* infection and/or bacteremia at the time of death in 7/8 Adélie penguins (Supplementary Table 3).

**Figure 4.**
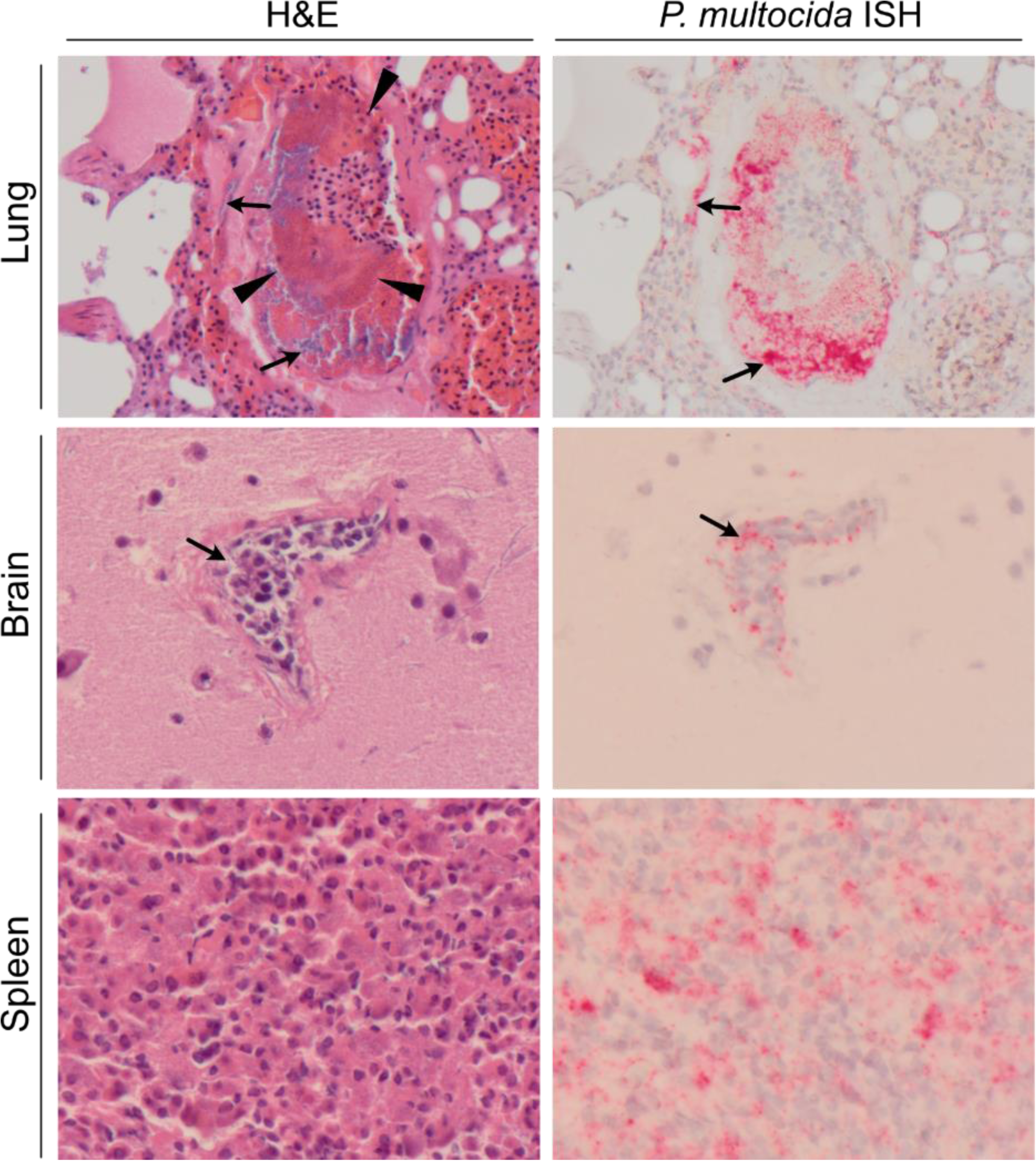
Serial sections of hematoxylin and eosin staining (H&E) and *P. multocida in situ* hybridization (ISH) of variable tissues of Adélie penguin. Lung, brain and spleen of Adelie penguin (AP50) found dead. Middle-sized to large blood vessels in lung and brain contain numerous small (1–2 µm diameter) coccoid bacteria in aggregates characteristic for *P. multocida*, staining positive (pink) in the ISH (arrows). In the lung, a larger blood vessel is distended by an aggregate of erythrocytes with pale cytoplasm, surrounded by strongly eosinophilic material that contains aggregates of bacteria (suspect thrombus, area with thrombus indicated with arrowheads). In the spleen, the distribution of bacteria is almost diffuse between the cells and less recognizable in the H&E.

In general, the semi-quantitative scoring of coccoid bacteria density in tissue slides stained with H&E matched well with those stained by *P. multocida-*specific ISH for Adélie penguins (Supplementary Table 3). The exceptions were spleens, in which the bacteria were abundantly observed in the ISH and not as clearly recognizable in the H&E as in other tissues. In the ISH-stained spleens, bacterial DNA was not observed in aggregates in middle to large blood vessels like in other tissues, but rather dispersed throughout lymphoid tissue. In addition, high *P. multocida* loads tested by qPCR (‘++’ and ‘+++’) matched well with the presence of lesions and *Pasteurella-*like coccoid bacteria within liver and/or blood vessels outside the liver (Supplementary Table 3).

Microscopic analyses did not corroborate *P. multocida* infection in the examined snowy sheathbill (SB02). *Pasteurella*-like bacteria were not observed in any of the tissues available for the snowy sheathbill, and all tissues were negative for *P. multocida* DNA and ISH staining (Supplementary Table 3).

#### IAV antigen was not detected in any tissues by immunohistochemistry

IAV NP antigen was not detected in any of the collected tissues, including those of two Adélie penguin carcasses in which low loads of IAV RNA were detected by RT-qPCR (Tables 1 and 2).

#### Avian cholera caused the death of most examined animals

Although IAV RNA was detected by RT-qPCR in 5/33 of the Adélie penguins, the low viral RNA loads, absence of influenza-like histologic lesions, and absence of IAV antigen expression in the tissues of the two of five penguins in which these tests were possible did not corroborate HPAI as the cause of death of these animals. Based on the combination of macroscopy, microscopy, ancillary testing for IAV and avian cholera, avian cholera was considered the most likely cause of death for 10/11 adult Adélie penguins at Heroina Island and 12/12 adult Adélie penguins at Beagle Island (Table 1). The causes of death were not as evident for the Adélie penguin chicks, with variable factors such as starvation, avian cholera, IAV infection, or a combination of these as possible contributors to the death of these animals (Table 1).

Several of the other animals found deceased were also considered likely to have died due to avian cholera (Table 1). Tissues from 7/8 skuas, 2/3 snowy sheathbills (SB03 and SB04) and 1/1 Antarctic fur seal (FS04) at Heroina Island and the 1/1 skua at Beagle Island had medium to high *P. multocida* DNA loads; however, poor tissue preservation obliterated histologic examination to determine the presence of characteristic lesions. Three of the skuas from Heroina Island (SK19, SK23 and SK24) also had low IAV RNA loads in the brain or lung, suggesting their deaths may have resulted from the combined effects of avian cholera and IAV. A snowy sheathbill (SB02) and the southern giant petrel (GP01) from Heroina Island had low *P. multocida* DNA loads, and their cause of death remains unclear.

#### Pasteurella multocida typing and phylogeny

Ten tissue samples (either liver, lung, or brain) from the Danger Islands with high *P. multocida* DNA loads (*kmt1* qPCR) were selected for sequencing. Of these, one sample from a snowy sheathbill (SB03 from Heroina Island) and five samples from Adélie penguins (AP38 and AP39 from Heroina Island; AP52, AP57, and AP58 from Beagle Island) had a *P. multocida* chromosome completeness ≥75%. Additionally, samples from another three seabirds collected at other Antarctic sites during the HPAI Australis Expedition were evaluated: liver samples from two skuas from Hope Bay (BSK01 and SK06) and a lung sample from one skua from Paulet Island (SK29). Multi-locus sequence typing revealed that for all samples the nearest allelic profile corresponded to the sequence type (ST) 61 in the RIRDC scheme (ST61; *adk4*, *est24*, *gdh2*, *mdh11*, *pgi21*, *pmi25*, *zwf20*), and ST91 in the multi-host scheme (*adk10, aroA33, deoD19, g6pd25, gdhA6, mdh12, pgi32*). Genomic serotyping revealed all samples had the exact or nearest capsular type A and LPS group 1 (A:L1; including the LPS Heddleston types 1 and 14) (Supplementary Table 4). The *Pasteurella multocida* toxin (*PMT*) gene was not present in any of the genomic sequences.

By mapping the *P. multocida* reads against the reference sequences, we obtained seven sequences with chromosome coverage above 82.5%, which were used for phylogenetic analysis. These were from skuas from Hope Bay (BSK01 and SK06) and Paulet Island (SK29), from a snowy sheathbill from Heroina Island (SB03), and from Adélie penguins from Heroina Island (AP38 and AP39) and Beagle Island (AP57). All these sequences belonged to a single phylogenetic lineage, and clustered together with sequences from seabirds sampled in 2019 at Amsterdam Island, Southern Indian Ocean (Figure 5, Supplementary Figure 2). The average nucleotide identity (ANI) among strains from the different geographic locations visited during our expedition ranged between 99.59%–99.86%, and ANI between these strains and the two Southern Indian Ocean reference genomes (CP097610 and CP097612) ranged between 99.63%–99.86%.

**Figure 5.**
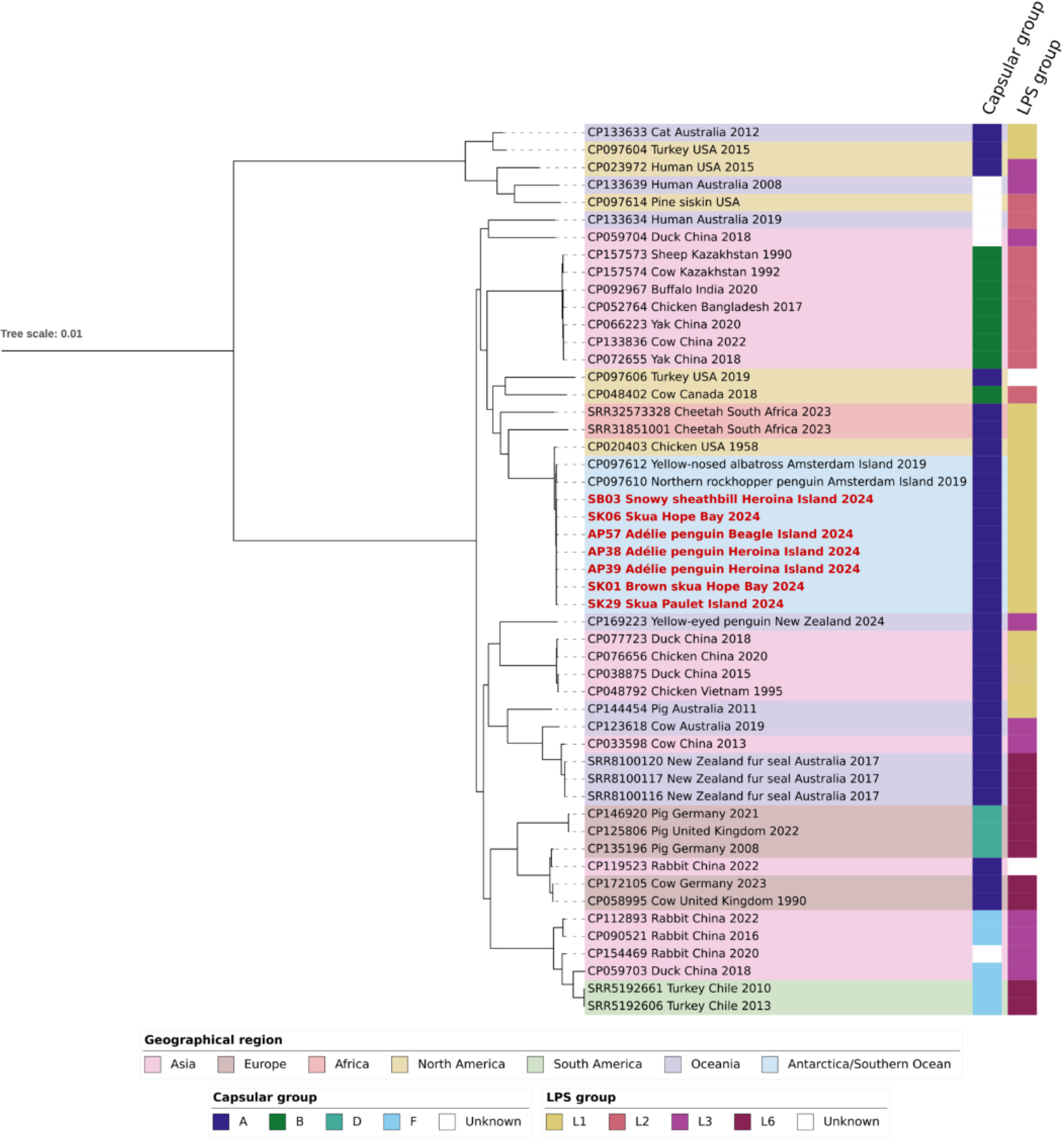
Phylogenetic analysis of *Pasteurella multocida* genomic sequences. Phylogenetic tree including *P. multocida* sequences from this study together with high quality sequences with available metadata on country of origin, collection date and host species. Capsular group, LPS type and geographical region are highlighted for all sequences. The extended version of this phylogenetic tree can be found in the supplementary materials.

## Discussion

During the HPAI Australis expedition in March 2024 we surveyed several sites across the Antarctic Peninsula and northern Weddell Sea [6,14], and came across a mass mortality event involving hundreds, possibly thousands, of birds at the Danger Islands. The most affected species was the Adélie penguin, but we also found carcasses of skuas, snowy sheathbills, southern giant petrel, and an Antarctic fur seal. Whereas mass mortalities of skuas at other sites in the Antarctic Peninsula in the summer of 2023/2024 were primarily attributed to H5N1 HPAIV of clade 2.3.4.4b [5,6], our investigations show that avian cholera played a primary role in the mortality event at the Danger Islands.

This interpretation was based on a combination of multiple diagnostic methods, including molecular analyses, microscopic analysis, and histological molecular localization techniques. Quantitative PCR targeting the *kmt1* gene of *P. multocida* yielded cycle threshold values indicative of heavy bacterial loads in multiple organs. Concurrent histopathological analyses of autopsied animals revealed hallmark lesions of avian cholera, and ISH demonstrated that these lesions were co-localized with detection of *P. multocida* DNA. In contrast, although low IAV RNA loads were detected in the tissues of some of the examined animals, there was neither IAV antigen nor influenza-like lesions. The combined findings, and the overall absence of other possible causes of death, provide robust evidence for the diagnosis of avian cholera in the examined animals.

During the incursion of H5N1 HPAIV into the Subantarctic and Antarctic regions in the 2023/2024 austral summer, we and others recorded sporadic or mass mortality in seabirds and pinnipeds linked to HPAIV at these locations, with skuas being significantly impacted [4–6,14,15]. While there was some evidence of IAV/HPAIV circulation at the Danger Islands both in this study and in previous testing [14,21], our multidisciplinary investigation does not corroborate HPAI as the cause of death. While it is possible that detectability of viral RNA was compromised by carcass degradation/autolysis, this seems unlikely given that we were able to detect moderate-high levels of host RNA (*GAPDH* housekeeping gene for seabirds and *β-actin* for seals) in the same samples. Our findings for the carcasses at the Danger Islands were also in sharp contrast with those of skuas sampled at other sites, where high HPAIV RNA loads were detected and virus antigen co-localized with lesions consistent with HPAIV [6]. Although HPAI was not the primary cause of mortality at the Danger Islands, it could have played a secondary role. This fits with our previous detections of HPAIV RNA in tissues from Adélie penguins and skuas that we diagnosed having a different cause of death [6,49]. The possibility of subclinical HPAIV infections in Adélie penguins would also be consistent with the detection of IAV H5 infections in apparently healthy penguins at Beagle Island and other sites in the Antarctic Peninsula [21].

Adélie penguins that we diagnosed with avian cholera showed the typical lesions associated with the disease, as described by others for Adélie penguins [3,6] and other seabirds [2,3,6,50,51]. Although we could only visit each island once, and did not observe sick Adélie penguins, the course of the disease was likely acute or peracute, based on the good nutritional condition of the carcasses and coagulative necrosis with influx of heterophils without a tissue repair response in their livers. Additionally, fecal swabs from healthy animals from Heroina Island, including Adélie penguins, tested negative for *P. multocida*, while cloacal swabs from carcasses of infected animals tested positive. These results, taken together, support the fast course of disease upon infection. It is difficult to estimate the chronology of the outbreak in an environment where carcasses were frozen. However, given the high density of penguin carcasses with a similar decomposition state it is possible that the epidemic was of relatively short duration. Since mass mortality of penguins was not witnessed during the visit to Heroina Island in late December 2023, and considering the breeding phenology of Adélie penguins, we estimate that the avian cholera outbreak likely occurred between January and early February 2024. Only one Antarctic fur seal was observed with non-specific clinical signs of disease, and although low HPAIV RNA loads were detected previously from another set of swabs from this animal [49], we did not detect IAV RNA here, and instead detected low *P. multocida* DNA loads. This difference is possibly related to the different biological materials in the secretions collected on different swab sets. Additionally, indirect collection of samples from this animal might have contributed to a negative IAV RNA result, and therefore neither disease can be ruled out.

Although our study marks the first report of an epidemic at the Danger Islands, it is possible that such epidemics have occurred before, considering the sporadic reports of avian cholera in the region since the late 1970s [2,6,36–41,50], but missed due to the lack of systematic annual monitoring. For comparison, there is almost annual recurrence of avian cholera outbreaks in water bird-concentrated wetlands [44,52], and in goose [53] and cormorant colonies in North America [54], and albatross colonies in the Southern Indian Ocean [38,40].

Phylogenetic analysis revealed that the *P. multocida* isolates detected in Adélie penguins and a sheathbill deceased at the Danger Islands clustered with isolates detected in deceased skuas at Hope Bay (∼120 km from Danger islands) and Paulet Island (∼60 km), from the same expedition. This suggests that a single strain has been circulating locally and caused seabird mortality in the Antarctic Peninsula and northern Weddell Sea. The pathogen could have been spread by subclinically infected animals. In previous studies, *P. multocida* with identical genetic profile had been recovered from a southern giant petrel carcass at Potter Peninsula (King George Island, South Shetland Islands) and Adélie penguin and skua carcasses at Hope Bay in the 1999/2000 austral summer, suggesting spread over a 150 km distance is possible [2,3].

The *P. multocida* genomic sequences from this study had the same sequence type (RIRDC ST61) and were phylogenetically closely related to *P. multocida* isolates recovered in 2019 from an Indian yellow-nosed albatross (*Thalassarche carteri*) and northern rockhopper penguins (*Eudyptes moseleyi*) at Amsterdam Island, Southern Indian Ocean, where recurring avian cholera outbreaks are a significant threat to seabird conservation since the mid-1980s [38,40,55–57]. Avian cholera has also been reported in other seabird communities in the Southern Ocean, including at Potter Peninsula [2], Palmer Station [36], Marion Island [37], and Campbell Island [50]. Interestingly, *P. multocida* strains recovered from a southern giant petrel at Potter Peninsula and from Adélie penguins at Hope Bay in 1999/2000 were of the capsular type A and LPS group 1 [2,3], like the one in our study. However, there is no genome sequencing data from the sites mentioned above for further comparison. It is noteworthy that the *P. multocida* isolates in this study and from seabirds in Amsterdam Island clustered with a *P. multocida* sequence recovered from a chicken in the United States in 1958. It was previously speculated that the *P. multocida* strain that has caused recurring outbreaks in seabirds at Amsterdam Island originated from poultry brought to Amsterdam Island in the early 1960s [38]. Our phylogenetic evidence goes further to suggest that the *P. multocida* isolates we detected in the Antarctic Peninsula showed high similarity with the *P. multocida* strain at Amsterdam Island. In contrast, the *P. multocida* isolate recovered from a yellow-eyed penguin (*Megadyptes antipodes*) at Otago, New Zealand [51], was not closely related to the isolates from this study or Amsterdam Island. This suggests that although long-distance dissemination of *P. multocida* among seabird populations in the Southern Ocean is plausible, the strains that infect Southern Ocean seabirds should not be presumed to be monophyletic. Our phylogenetic analysis has some limitations. Due to the remote sampling location and biosafety concerns, our samples were preserved in buffers to inactivate pathogens and stabilize nucleic acids. As a result, *P. multocida* could not be cultured for whole genome sequencing. Instead, reference-guided draft genome sequences were reconstructed from metagenomic data, which impacts genome completeness, quality and strain-level resolution and serotyping. Additionally, the lack of *P. multocida* sequencing data from other sites in the Subantarctic and Antarctic region limits our assessment of the most likely introduction and transmission patterns. Nevertheless, our results show the value of such analyses to determine the most likely pattern of introduction and spread.

The reservoirs and vectors of *P. multocida* in the Southern Ocean are currently unclear. In avian cholera outbreaks in waterfowl in North America, there is no evidence for persistence of *P. multocida* in wetland ecosystems where migratory waterfowl stage and winter [58]. Instead, a small proportion of snow geese (*Anser caerulescens*) and Ross’s geese (*Chen rossii*) are subclinical carriers of pathogenic variants of *P. multocida* and can act as the reservoir of avian cholera [59,60]. Interestingly, previous studies noted that kelp gulls with *P. multocida* at Hope Bay had lesions compatible with chronic avian cholera, and the isolates were only recovered from the mouth [3]. Furthermore, antibodies against *P. multocida* were detected in brown skuas and albatrosses at Amsterdam Island [55], suggesting these species may survive the infections and potentially serve as reservoirs or vectors for *P. multocida* dissemination. However, it remains to be determined whether Adélie penguins or other species could also be subclinical carriers of *P. multocida*, or alternatively, whether the low ambient temperatures present in Antarctica most of the year may allow the bacterium to persist in the environment of the Danger Islands over the winter. Since *P. multocida* has been shown to persist for up to 100–120 days in soil and water [44], the cold Antarctic environment, combined with the presence of frozen carcasses, soil and freshwater, could provide conditions for the bacterium to survive through the Antarctic winter.

Regarding transmission, investigation of avian cholera outbreaks in wild birds in North America suggests that both contaminated water and bird-to-bird contact can play key roles [44,61]. In California gulls (*Larus californicus*) fed contaminated meat, the bacterium was detected in feces for up to 5 days [44]. Although we did not systematically test cloacal swabs or intestine samples from Adélie penguin carcasses, in the small number of samples evaluated we did detect *P. multocida* DNA loads that were higher than those of brain and lung samples from the same individuals, suggesting that fecal shedding may be significant and fecal-oral transmission could have occurred. In the avian cholera outbreaks previously reported at Hope Bay, it was considered that freshwater lakes played a key role in the transmission of *P. multocida* to Adélie penguins [3]. Although such lakes are absent in the Danger Islands, snow and small puddles of water contaminated with penguin feces, could have played a similar role. The remarkable high density of Adélie penguins at both islands also provides plentiful opportunities for direct bird-to-bird transmission, such as exposure to aerosols and oronasal secretions from infected individuals. Collection of feces-contaminated rocks to build their nests could also provide opportunities for transmission. Transmission to predatory and scavenging species such as skuas and sheathbills most likely occurred through consumption of infected animals [56], as has been described for avian cholera cases in North America [46]. Additionally, for Adélie penguins, the occasional pecking and scavenging of other Adélie penguin carcasses [62], necrophilic behavior [63] and fighting [64] might have contributed to a lesser extent.

Although Adélie penguins are not currently threatened with extinction and their global population is increasing [65], in spite of a decrease in the Western side of the Antarctic Peninsula [28,66], the documented outbreak of avian cholera at the Danger Islands is concerning for several reasons. This site represents a sanctuary for the conservation of this species, hosting 7.5% of its global population and 55% of the Antarctic Peninsula population [24,32]. The repeated occurrence of avian cholera outbreaks in this population could therefore pose a significant threat on a regional scale, especially if the impacts of outbreaks synergize with other local threats such as HPAIV, climate change, and krill harvesting [27]. Additionally, the potential recurrence of avian cholera outbreaks in Adélie penguins on the Danger Islands could represent a source of infection and mortality for other avian and mammalian species in the region.

In conclusion, the discovery of avian cholera as a cause of seabird mass mortality in Antarctica concurrently with the expansion of HPAIV into this continent highlights the importance of considering this disease in the differential diagnosis of unusual mortality events in Antarctic wildlife. While targeted surveillance is essential to understand the spread and impacts of emerging pathogens such as HPAIV, a more comprehensive approach that considers differential diagnoses and incorporates detailed pathological investigation to be able to distinguish between different factors driving mortality can be invaluable to ensure that the epidemiological context and the relative roles of co-occurring pathogens are fully appreciated. Our findings underscore several knowledge gaps on the occurrence, epidemiology and impacts of avian cholera in Southern Ocean wildlife, and provide methodological tools to investigate this disease during wildlife mortality events. Understanding the potential host and environmental reservoirs of *P. multocida*, the mechanisms of local and global strain dissemination, the triggers of outbreaks in natural seabird populations, and the potential impacts on bird and mammal populations in the region is relevant to increase awareness of the risks posed by infectious diseases in Antarctica and, in the future, might also prompt policy frameworks that enable effective response and long-term management of this region under a One Health perspective.

## Materials and methods

### Site assessment and sample collection

All the samples described in this study have been collected between March 25^th^ and March 26^th^, 2024, at Heroina Island (63°24’0”S, 54°36’0”W) and Beagle Island (63°25′0″S, 54°40′0″W). Sample collection was undertaken as described previously [6]. Fresh oronasal and peri-anal secretions from the live Antarctic fur seal at Heroina Island (FS03) were swabbed from the snow and rocks where the animal was resting, without touching the animal. Samples for virological and bacteriological analyses and for histopathology were collected based on two main objectives: a) determine HPAI impact on the species found dead (which was the initial aim of the expedition), and b) collecting samples from as many carcasses as possible for diagnostics purposes. Based on these two objectives, we collected three different types of sample sets:

1. Full set: this includes samples for virological and bacteriological analyses and samples for histopathological analyses. Sampled organs are trachea, brain, lung, heart, liver, intestine, spleen, and kidney. Oropharyngeal and cloacal swabs and feather follicles were also collected. Whether all or a subset of these samples were collected was based on the state of autolysis of each animal. The number of carcasses sampled this way was limited by time and by the number of carcasses present at each site. Three carcasses (AP37, AP38, AP48) are included in this category.
2. Medium set: this includes brain and/or lung tissues for virological and bacteriological analyses and variable samples for histopathological analyses as described in 1), based on autolysis and scavenging state of the carcasses. Six carcasses (AP27, AP29, AP33, AP50, AP51, SB02) are included in this category.
3. Minimum set: this includes brain and/or lung swabs for virological and bacteriological analyses, collected without any macroscopic observation of the organs. All the carcasses not included in the two previous categories are included here.

At Heroina Island, a comprehensive search for carcasses and symptomatic animals was conducted on the western peninsula and the main landing ramp (approximately 22,000 m^2^; red area shown in Figure 1C), with carcasses being examined on an individual basis. A team also climbed the northern plateau of Heroina Island to conduct a superficial assessment of the extent of mortality; the island’s southern plateau was not visited. Time ashore at Beagle Island was extremely limited (1.5 hours) due to snowfall and the need to move forward for navigation conditions, hence only a superficial assessment of the mortality could be conducted. Species were identified based on external characteristics [23]; any individual with down feathers was classified as a chick. The levels of confidence of diagnoses are described in Supplementary Table 5. Personal protective equipment and permits for sample collection were the same as described previously [6].

### Nucleic acid extraction and qPCRs

Nucleic acid from animal tissues and swabs for virological and bacteriological analyses, qPCRs for *M*, *H5*, *N1*, HPAI, *kmt1*, *GAPDH* and *β-actin*, and Sanger sequencing of the MBCS and the *kmt1* amplicon were performed as described previously [6]. Fecal swab supernatants were spun down at 13,000 g for 5 minutes and diluted 1:5 in PBS. Then, 200 µl of each diluted supernatant was added to 600 µl of MagNA Pure 96 External Lysis Buffer (Roche) and PBS up to 1 ml; 10 µl of phocine distemper virus was added to each sample [67] as extraction control and PCR inhibition control. Total nucleic acids were extracted using MagNA Pure 96 System (Roche). The methodology for RNA extraction from fecal samples and IAV RT-qPCR is the same as described previously [6]. The following categories were established RNA/DNA load (40 – Cq): 0, negative (-); 1 to 8, low (+); 9 to 16, moderate (++); 17 to 40, high (+++).

### Metagenomic sequencing and sequence data analysis

Nucleic acids were extracted using the High Pure Nucleic Acid Kit (Roche). DNA was then randomly amplified using Sequence-independent single-primer amplification (SISPA) [68] followed by sequencing library preparation from 100 ng of product using the Native Barcoding Kit 24 v14 (Oxford Nanopore Technologies). Eighteen samples and a non-template control were run on a FLO-PRO114M flow cell on a PromethION (Oxford Nanopore Technologies). Sequencing reads were basecalled using the super-accurate basecalling model and barcodes were trimmed in Dorado Basecall Server 7.4.12 (Oxford Nanopore Technologies). SISPA primer sequences were removed using Cutadapt v3.0 (https://github.com/marcelm/cutadapt) and reads trimmed with fastq v0.25 (https://github.com/mcollina/fastq). Host reads were removed using minimap2 v2.28 (https://github.com/lh3/minimap2). Reads were mapped against 622 *Pasteurella multocida* RefSeq sequences (accessed 11/02/2025, https://www.ncbi.nlm.nih.gov/nuccore) to find the closest reference based on mapped reads and coverage using Samtools v1.22 (depth) (https://github.com/samtools) and Bedtools v2.31 (genomecov) (https://github.com/arq5x/bedtools2). Draft genomic consensus sequences were generated using Virconsens pipeline (https://github.com/dnieuw/Virconsens). Plasmid reads were not analysed. The sequence type (ST) was identified using MLST Finder v2.0 using the *Pasteurella* RIRDC MLST scheme [69]. Capsular types and LPS groups were identified using the database for whole genomic prediction of serotypes developed by the Department of Veterinary and Animal Sciences of the University of Copenhagen (https://ivsmlst.sund.ku.dk/) [70].

For phylogenetic analysis, high quality sequences with available metadata on country of origin, collection date and host species were downloaded from the bacterial and viral bioinformatics resource center (accessed 16/04/2025, https://www.bv-brc.org/). Sequences above 80% coverage from our sample collection were arbitrarily included in the phylogenetic analysis. Sequences were aligned with minimap2 (https://github.com/lh3/minimap2) followed by gofasta (https://github.com/virus-evolution/gofasta) and recombinant sequences were removed with gubbins v3.4 (https://github.com/nickjcroucher/gubbins). IQ-TREE v2.4.0 was used with ModelFinder and 1000 ultrafast bootstraps [71]. Average nucleotide identity (ANI) was calculated using ANIb (BLAST+).

### Immunohistochemistry, *in situ* hybridization, and histopathology

IHC for the NP antigen of influenza A virus was performed as described previously [6]. ISH for *Pasteurella multocida* DNA was performed as described previously, including the positive controls [6]. Formalin-fixed paraffin-embedded tissue sections (3 µm) were deparaffinized using xylene, rehydrated using graded ethanol, stained with hematoxylin (Klinipath) and eosin (QPath; H&E), and assessed by use of a light microscope by a qualified veterinary pathologist (LB) for any histopathological changes.

## Acknowledgments

We are grateful to Antonio Alcamí and Begoña Aguado for their support and contribution as part of the HPAI Australis Expedition team. We thank Bas Oude Munnink for support with the *Pasteurella multocida* sequencing and phylogenetic analysis, Timm Harder, Sanne Thewessen and Ron Fouchier for support in the molecular analyses, Jurriaan de Steenwinkel for providing a *P. multocida* positive control, Michelle Wille and Gabriele Arcari for the preliminary consultation for bacterial metagenomics.

## Author contributions

Conceptualization: MI, AG, FS, MB, MD, TK, LB, RETV

Methodology: MI, AG, LS, LL, TMB, SL, LB, RETV

Investigation: MI, AG, LS, FS, LL, TMB, SL, MD, LB, RETV

Visualization: MI, AG, TK, LB, RETV

Sailing activities: AC, AR, BW

Writing – original draft: MI, TK, LB, RETV

Writing – review & editing: all authors

