## Supplementary materials for "Mass mortality at penguin mega-colonies due to avian cholera confounds H5N1 HPAIV surveillance in Antarctica"

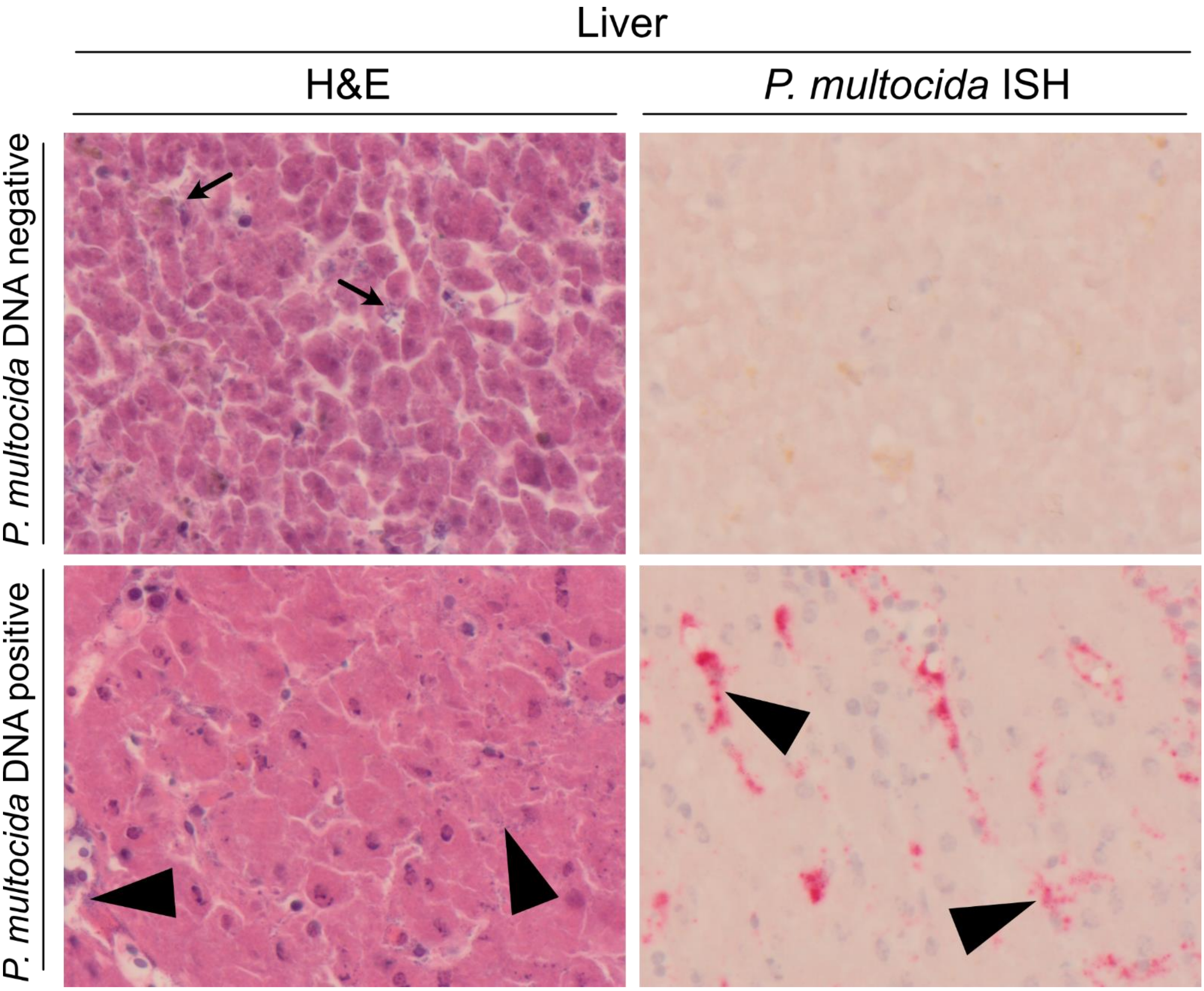


**Supplementary Figure 1. *P. multocida* qPCR-negative and qPCR-positive liver of Adélie penguin.**
Serial sections of hematoxylin and eosin staining (H&E) and *P. multocida in situ* hybridization (ISH) in the liver of Adélie penguins. Livers of Adélie penguins found dead with similar levels of autolysis are shown, with a *P. multocida* qPCR-negative individual (AP03) compared to a *P. multocida* qPCR-positive individual (AP50). The qPCR-negative liver shows a variety of rod-shaped and coccoid bacteria (arrows) dispersed between hepatocytes, characteristic of postmortem overgrowth, and no staining by *P. multocida* ISH. The qPCR-positive liver shows small coccoid bacteria within sinusoids, and brightly pink positive staining by *P. multocida* ISH within sinusoids (arrowheads).

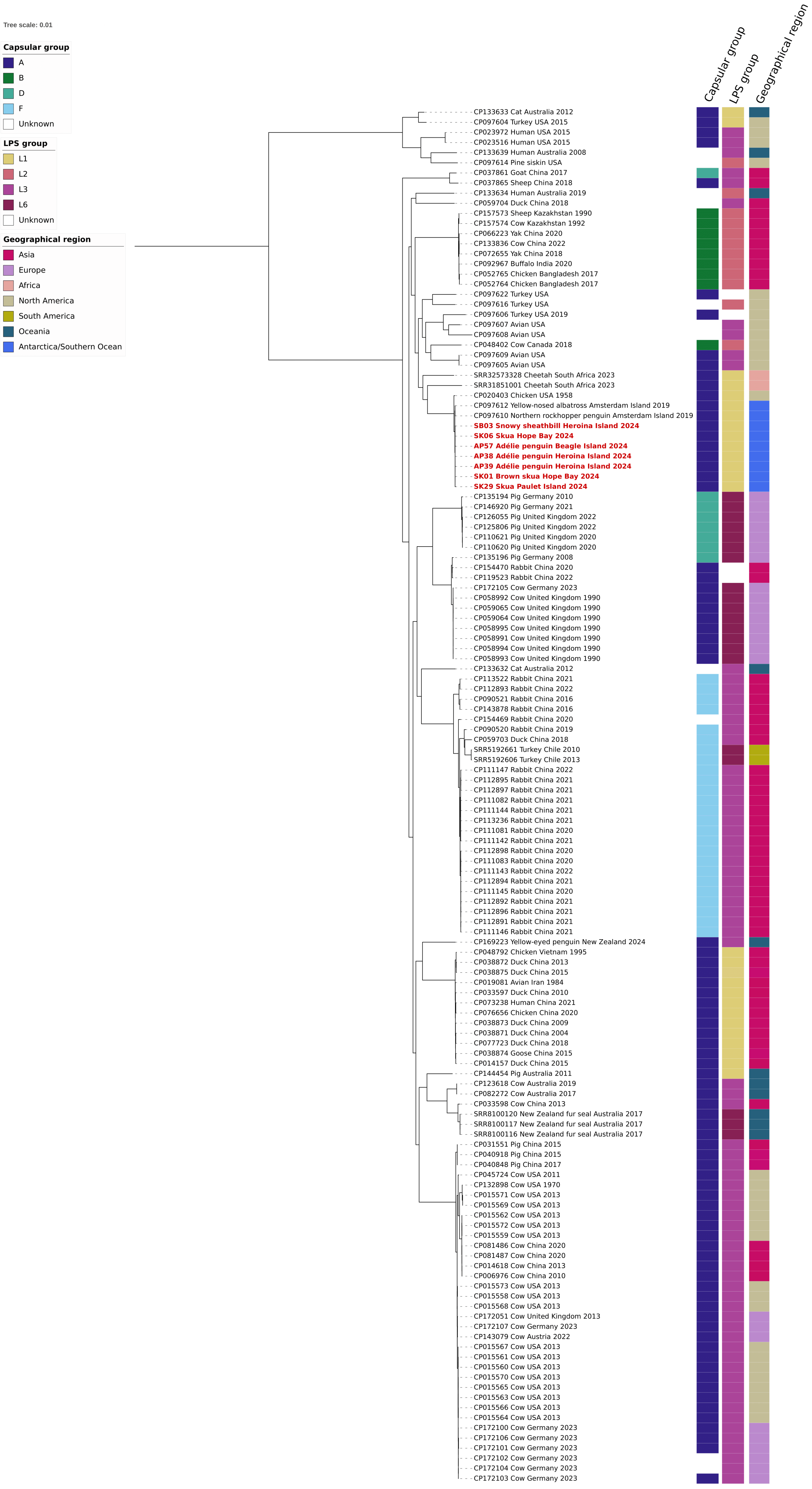


**Supplementary Figure 2. Extended phylogenetic analysis of *Pasteurella multocida* genomic sequences.** Phylogenetic tree including *P. multocida* sequences from this study together with high quality sequences with available metadata on country of origin, collection date and host species. Capsular group, LPS type and geographical region are highlighted for all sequences.

**Supplementary table 1. Results of influenza A virus RT-qPCRs (*M, H5, N1* ; blue) and *P. multocida* qPCR (*kmt1* ; red) of swabs and tissues from animals found dead from Heroina and Beagle Island.** Results are shown as Cq values. Since all samples tested negative in the HPAI RT-qPCR, the results from this RT-qPCR are not shown. The IDs from onboard testing are indicated in brackets to compare with previous results (14, 49).

| Location | Species | ID | ID onboard test (49) | Oropharyngeal swab | | | | Cloacal swab | | | | Brain | | | | Lung | | | | Intestine | | | | Liver | | | | Spleen | | | | Kidney | | | | Feather follicle | | | |
|---|---|---|---|---|---|---|---|---|---|---|---|---|---|---|---|---|---|---|---|---|---|---|---|---|---|---|---|---|---|---|---|---|---|---|---|---|---|---|---|
| | | | | M1 | H5 | N1 | kmt1 | M1 | H5 | N1 | kmt1 | M1 | H5 | N1 | kmt1 | M1 | H5 | N1 | kmt1 | M1 | H5 | N1 | kmt1 | M1 | H5 | N1 | kmt1 | M1 | H5 | N1 | kmt1 | M1 | H5 | N1 | kmt1 | M1 | H5 | N1 | kmt1 |
| **Heroina Island** | **Adélie penguin** (*Pygoscelis* | **AP27** | ADPE_1_Hero | na | na | na | na | na | na | na | na | nd | nd | na | 22.8 | nd | nd | na | 21.3 | na | na | na | na | na | na | na | na | na | na | na | na | na | na | na | na | na | na | na | na |
| | *adeliae* ) | **AP28** | ADPE_2_Hero | na | na | na | na | na | na | na | na | nd | nd | na | 19.7 | nd | nd | na | 20.1 | na | na | na | na | na | na | na | na | na | na | na | na | na | na | na | na | na | na | na | na |
| | | **AP29** | ADPE_3_Hero | na | na | na | na | na | na | na | na | nd | nd | na | 20.9 | nd | nd | na | 19.8 | na | na | na | na | na | na | na | na | na | na | na | na | na | na | na | na | na | na | na | na |
| | | **AP30** | | na | na | na | na | na | na | na | na | nd | nd | na | 20.8 | nd | nd | na | 19.8 | na | na | na | na | na | na | na | na | na | na | na | na | na | na | na | na | na | na | na | na |
| | | **AP31** | | na | na | na | na | na | na | na | na | nd | nd | na | 22.6 | 32.5 | nd | na | 21.4 | na | na | na | na | na | na | na | na | na | na | na | na | na | na | na | na | na | na | na | na |
| | | **AP32** | | na | na | na | na | na | na | na | na | nd | nd | na | 21.0 | nd | nd | na | 19.0 | na | na | na | na | na | na | na | na | na | na | na | na | na | na | na | na | na | na | na | na |
| | | **AP33** | | na | na | na | na | na | na | na | na | nd | nd | na | 29.3 | nd | nd | na | 27.0 | na | na | na | na | na | na | na | na | na | na | na | na | na | na | na | na | na | na | na | na |
| | | **AP34** | | na | na | na | na | na | na | na | na | nd | nd | na | 29.1 | nd | nd | na | 29.7 | na | na | na | na | na | na | na | na | na | na | na | na | na | na | na | na | na | na | na | na |
| | | **AP35** | ADPE_4_Hero | na | na | na | na | na | na | na | na | nd | nd | na | 31.6 | nd | nd | na | 30.0 | na | na | na | na | na | na | na | na | na | na | na | na | na | na | na | na | na | na | na | na |
| | | **AP36** | | na | na | na | na | na | na | na | na | nd | nd | na | 33.0 | nd | nd | na | 30.0 | na | na | na | na | na | na | na | na | na | na | na | na | na | na | na | na | na | na | na | na |
| | | **AP37** | ADPE_8_Hero | 33.6 | nd | na | 31.4 | nd | nd | na | 29.8 | nd | nd | na | 34.5 | nd | nd | na | 31.4 | 35.3 | nd | na | 29.5 | 32.1 | 34.2 | 36.4 | 28.2 | nd | nd | na | 36.2 | nd | nd | na | 36.1 | 32.7 | 33.9 | na | 29.3 |
| | | **AP38** | | nd | nd | na | 25.9 | na | na | na | na | nd | nd | na | 19.4 | nd | nd | na | 18.7 | 35.1 | nd | na | 16.6 | 34.7 | nd | na | 17.4 | na | na | na | na | nd | nd | na | 19.6 | 34.8 | nd | na | 25.7 |
| | | **AP39** | ADPE_9_Hero | na | na | na | na | na | na | na | na | nd | nd | na | 19.7 | nd | nd | na | 18.5 | na | na | na | na | na | na | na | na | na | na | na | na | na | na | na | na | na | na | na | na |
| | | **AP40** | ADPE_10_Hero | na | na | na | na | na | na | na | na | nd | nd | na | 20.9 | nd | nd | na | 20.0 | na | na | na | na | na | na | na | na | na | na | na | na | na | na | na | na | na | na | na | na |
| | | **AP41** | ADPE_11_Hero | na | na | na | na | na | na | na | na | nd | nd | na | 20.3 | 36.6 | nd | na | 20.5 | na | na | na | na | na | na | na | na | na | na | na | na | na | na | na | na | na | na | na | na |
| | | **AP42** | | na | na | na | na | na | na | na | na | nd | nd | na | 35.5 | nd | nd | na | 36.1 | na | na | na | na | na | na | na | na | na | na | na | na | na | na | na | na | na | na | na | na |
| | | **AP43** | ADPE_5_Hero | na | na | na | na | na | na | na | na | nd | nd | na | nd | nd | nd | na | 35.9 | na | na | na | na | na | na | na | na | na | na | na | na | na | na | na | na | na | na | na | na |
| | | **AP44** | | na | na | na | na | na | na | na | na | nd | nd | na | nd | nd | nd | na | nd | na | na | na | na | na | na | na | na | na | na | na | na | na | na | na | na | na | na | na | na |
| | | **AP45** | | na | na | na | na | na | na | na | na | nd | nd | na | nd | nd | nd | na | nd | na | na | na | na | na | na | na | na | na | na | na | na | na | na | na | na | na | na | na | na |
| | | **AP46** | ADPE_6_Hero | na | na | na | na | na | na | na | na | nd | nd | na | nd | nd | nd | na | nd | na | na | na | na | na | na | na | na | na | na | na | na | na | na | na | na | na | na | na | na |
| | | **AP47** | ADPE_7_Hero | na | na | na | na | na | na | na | na | 32.3 | 33.2 | nd | nd | 34.3 | 35.8 | nd | 36.4 | na | na | na | na | na | na | na | na | na | na | na | na | na | na | na | na | na | na | na | na |
| | **Skua** | **SK18** | Skua_1_Hero | na | na | na | na | na | na | na | na | nd | nd | na | 21.0 | nd | nd | na | 17.7 | na | na | na | na | na | na | na | na | na | na | na | na | na | na | na | na | na | na | na | na |
| | (*Stercorarius* sp.) | **SK19** | Skua_2_Hero | na | na | na | na | na | na | na | na | 36.4 | nd | na | 25.0 | nd | nd | na | 19.4 | na | na | na | na | na | na | na | na | na | na | na | na | na | na | na | na | na | na | na | na |
| | | **BSK20** | Skua_3_Hero | na | na | na | na | na | na | na | na | nd | nd | na | 23.3 | nd | nd | na | 18.0 | na | na | na | na | na | na | na | na | na | na | na | na | na | na | na | na | na | na | na | na |
| | | **BSK21** | | na | na | na | na | na | na | na | na | nd | nd | na | 24.8 | na | na | na | na | na | na | na | na | na | na | na | na | na | na | na | na | na | na | na | na | na | na | na | na |
| | | **SK22** | | na | na | na | na | na | na | na | na | nd | nd | na | 22.7 | na | na | na | na | na | na | na | na | na | na | na | na | na | na | na | na | na | na | na | na | na | na | na | na |
| | | **SK23** | | na | na | na | na | na | na | na | na | 34,5 | nd | na | 27.1 | na | na | na | na | na | na | na | na | na | na | na | na | na | na | na | na | na | na | na | na | na | na | na | na |
| | | **SK24** | Skua_4_Hero | na | na | na | na | na | na | na | na | nd | nd | na | 32.9 | 33.6 | nd | na | 22.1 | na | na | na | na | na | na | na | na | na | na | na | na | na | na | na | na | na | na | na | na |
| | | **BSK25** | | na | na | na | na | na | na | na | na | nd | nd | na | 27.3 | na | na | na | na | na | na | na | na | na | na | na | na | na | na | na | na | na | na | na | na | na | na | na | na |
| | | **SK26** | | na | na | na | na | na | na | na | na | nd | nd | na | 28.4 | na | na | na | na | na | na | na | na | na | na | na | na | na | na | na | na | na | na | na | na | na | na | na | na |
| | | **SK27** | | na | na | na | na | na | na | na | na | nd | nd | na | 29.2 | na | na | na | na | na | na | na | na | na | na | na | na | na | na | na | na | na | na | na | na | na | na | na | na |
| | **Snowy sheathbill** | **SB02** | SSB_3_Hero | na | na | na | na | na | na | na | na | nd | nd | na | 35.6 | nd | nd | na | 28.1 | na | na | na | na | na | na | na | na | na | na | na | na | na | na | na | na | na | na | na | na |
| | (*Chionis albus* ) | **SB03** | SSB_1_Hero | na | na | na | na | na | na | na | na | nd | nd | na | 21.1 | na | na | na | na | na | na | na | na | na | na | na | na | na | na | na | na | na | na | na | na | na | na | na | na |
| | | **SB04** | SSB_2_Hero | na | na | na | na | na | na | na | na | nd | nd | na | 28.9 | na | na | na | na | na | na | na | na | na | na | na | na | na | na | na | na | na | na | na | na | na | na | na | na |
| | **Southern giant petrel** (*Macronectes giganteus* ) | **GP01** | GP_1_Hero | na | na | na | na | na | na | na | na | nd | nd | na | 33.4 | na | na | na | na | na | na | na | na | na | na | na | na | na | na | na | na | na | na | na | na | na | na | na | na |
| | **Antarctic fur seal** | **FS03.1*** | AFS_alive_2_Hero | nd# | nd# | na | 34.1 | nd# | nd# | na | nd | na | na | na | na | na | na | na | na | na | na | na | na | na | na | na | na | na | na | na | na | na | na | na | na | na | na | na | na |
| | (*Arctocephalus gazella* ) | **FS03.2*** | AFS_alive_2_Hero | nd# | nd# | na | nd | nd# | nd# | na | 36.8 | na | na | na | na | na | na | na | na | na | na | na | na | na | na | na | na | na | na | na | na | na | na | na | na | na | na | na | na |
| | | **FS04** | AFS_1_Hero | na | na | na | na | na | na | na | na | nd | nd | na | 25.8 | na | na | na | na | na | na | na | na | na | na | na | na | na | na | na | na | na | na | na | na | na | na | na | na |
| **Beagle Island** | **Adélie penguin** (*Pygosce lis* | **AP48** | ADPE_1_Beagle | nd | nd | na | 24.7 | nd | nd | na | 27.0 | nd | nd | na | 18.2 | nd | nd | na | 20.4 | nd | nd | na | 23.0 | nd | nd | na | 16.2 | na | na | na | na | nd | nd | na | 18.2 | nd | nd | na | 23.4 |
| | *adeliae* ) | **AP49** | ADPE_2_Beagle | na | na | na | na | na | na | na | na | nd | nd | na | 20.3 | nd | nd | na | 21.4 | na | na | na | na | na | na | na | na | na | na | na | na | na | na | na | na | na | na | na | na |
| | | **AP50** | ADPE_3_Beagle | na | na | na | na | na | na | na | na | nd | nd | na | 21.8 | nd | nd | na | 18.1 | na | na | na | na | na | na | na | na | na | na | na | na | na | na | na | na | na | na | na | na |
| | | **AP51** | ADPE_4_Beagle | na | na | na | na | na | na | na | na | nd | nd | na | 19.4 | nd | nd | na | 17.7 | na | na | na | na | na | na | na | na | na | na | na | na | na | na | na | na | na | na | na | na |
| | | **AP52** | ADPE_5_Beagle | na | na | na | na | na | na | na | na | nd | nd | na | 18.7 | nd | nd | na | 18.7 | na | na | na | na | na | na | na | na | na | na | na | na | na | na | na | na | na | na | na | na |
| | | **AP53** | ADPE_6_Beagle | na | na | na | na | na | na | na | na | nd | nd | na | 19.9 | nd | nd | na | 19.0 | na | na | na | na | na | na | na | na | na | na | na | na | na | na | na | na | na | na | na | na |
| | | **AP54** | ADPE_7_Beagle | na | na | na | na | na | na | na | na | nd | nd | na | 27.4 | nd | nd | na | 26.5 | na | na | na | na | na | na | na | na | na | na | na | na | na | na | na | na | na | na | na | na |
| | | **AP55** | ADPE_8_Beagle | na | na | na | na | na | na | na | na | nd | nd | na | 20.3 | na | na | na | na | na | na | na | na | na | na | na | na | na | na | na | na | na | na | na | na | na | na | na | na |
| | | **AP56** | ADPE_9_Beagle | na | na | na | na | na | na | na | na | nd | nd | na | 19.3 | na | na | na | na | na | na | na | na | na | na | na | na | na | na | na | na | na | na | na | na | na | na | na | na |
| | | **AP57** | ADPE_10_Beagle | na | na | na | na | na | na | na | na | nd | nd | na | 18.5 | na | na | na | na | na | na | na | na | na | na | na | na | na | na | na | na | na | na | na | na | na | na | na | na |
| | | **AP58** | ADPE_11_Beagle | na | na | na | na | na | na | na | na | nd | nd | na | 20.6 | nd | nd | na | 18.2 | na | na | na | na | na | na | na | na | na | na | na | na | na | na | na | na | na | na | na | na |
| | | **AP59** | ADPE_12_Beagle | na | na | na | na | na | na | na | na | nd | nd | na | 19.2 | na | na | na | na | na | na | na | na | na | na | na | na | na | na | na | na | na | na | na | na | na | na | na | na |
| | **Skua** (*Stercorarius* sp.) | **SK28** | Skua_1_Beagle | na | na | na | na | na | na | na | na | nd | nd | na | 27.5 | nd | nd | na | 24.1 | na | na | na | na | na | na | na | na | na | na | na | na | na | na | na | na | na | na | na | na |

* FS03 was alive and environmental oro-nasal and peri-anal secretions were sampled in two consecutive days.

### These values refer to environmental oro-nasal secretion swab (indicated as oropharyngeal swab) and peri-anal secretion swab (indicated as cloacal swab).

na: not applicable (not sampled)

nd: not detected

**Supplementary table 2. Housekeeping gene RNA loads (40-Cq) in swabs (s.) and tissues of animals found dead at Heroina and Beagle Island.** GAPDH RNA is targeted for birds and β-actin RNA for seals.

| Location | Species | ID | Oropharyngeal s. | Cloacal s. | Brain | Lung | Intestine | Liver | Spleen | Kidney | Feather follicle |
|---|---|---|---|---|---|---|---|---|---|---|---|
| **Heroina Island** | **Adélie penguin** (*Pygoscelis adeliae* ) | **AP27** | na | na | 19,5 | 21,0 | na | na | na | na | na |
| | | **AP28** | na | na | 21,0 | 21,4 | na | na | na | na | na |
| | | **AP29** | na | na | 20,4 | 22,6 | na | na | na | na | na |
| | | **AP30** | na | na | 19,4 | 22,0 | na | na | na | na | na |
| | | **AP31** | na | na | 22,6 | 21,2 | na | na | na | na | na |
| | | **AP32** | na | na | 19,8 | 21,7 | na | na | na | na | na |
| | | **AP33** | na | na | 19,3 | 20,9 | na | na | na | na | na |
| | | **AP34** | na | na | 20,8 | 21,1 | na | na | na | na | na |
| | | **AP35** | na | na | 20,9 | 20,9 | na | na | na | na | na |
| | | **AP36** | na | na | 20,0 | 20,9 | na | na | na | na | na |
| | | **AP37** | 12,8 | 14,6 | 20,0 | 21,0 | 20,5 | 19,3 | 22,1 | 24,0 | 22,3 |
| | | **AP38** | 18,0 | na | 19,4 | 21,1 | 19,8 | 20,5 | na | 22,6 | 18,8 |
| | | **AP39** | na | na | 21,3 | 21,4 | na | na | na | na | na |
| | | **AP40** | na | na | 20,7 | 22,2 | na | na | na | na | na |
| | | **AP41** | na | na | 20,8 | 22,6 | na | na | na | na | na |
| | | **AP42** | na | na | 20,5 | 21,6 | na | na | na | na | na |
| | | **AP43** | na | na | 21,9 | 22,1 | na | na | na | na | na |
| | | **AP44** | na | na | 20,7 | 22,0 | na | na | na | na | na |
| | | **AP45** | na | na | 21,1 | 21,8 | na | na | na | na | na |
| | | **AP46** | na | na | 21,4 | 21,2 | na | na | na | na | na |
| | | **AP47** | na | na | 21,5 | 20,4 | na | na | na | na | na |
| | **Skua** (*Stercorarius* sp.) | **SK18** | na | na | 21,9 | 18,5 | na | na | na | na | na |
| | | **SK19** | na | na | 18,8 | 16,4 | na | na | na | na | na |
| | | **BSK20** | na | na | 19,3 | 20,7 | na | na | na | na | na |
| | | **BSK21** | na | na | 21,0 | na | na | na | na | na | na |
| | | **SK22** | na | na | 21,1 | na | na | na | na | na | na |
| | | **SK23** | na | na | 19,9 | na | na | na | na | na | na |
| | | **SK24** | na | na | 21,3 | 21,6 | na | na | na | na | na |
| | | **BSK25** | na | na | 21,0 | na | na | na | na | na | na |
| | | **SK26** | na | na | 19,8 | na | na | na | na | na | na |
| | | **SK27** | na | na | 19,7 | na | na | na | na | na | na |
| | **Snowy sheathbill** (*Chionis albus* ) | **SB02** | na | na | 21,5 | 22,5 | na | na | na | na | na |
| | | **SB03** | na | na | 19,2 | na | na | na | na | na | na |
| | | **SB04** | na | na | 21,9 | na | na | na | na | na | na |
| | **Southern giant petrel** (*Macronectes giganteus* ) | **GP01** | na | na | 19,4 | na | na | na | na | na | na |
| | **Antarctic fur seal** (*Arctocephalus gazella* ) | **FS03.1*** | 13.1# | 6.8# | na | na | na | na | na | na | na |
| | | **FS03.2*** | nd# | nd# | na | na | na | na | na | na | na |
| | | **FS04** | na | na | 16,4 | na | na | na | na | na | na |
| **Beagle Island** | **Adélie penguin** *(Pygosce* lis adeliae) | **AP48** | 16,3 | 17,5 | 21,3 | 22,6 | 22,9 | 20,0 | na | 22,4 | 20,5 |
| | | **AP49** | na | na | 19,8 | 19,5 | na | na | na | na | na |
| | | **AP50** | na | na | 18,3 | 19,3 | na | na | na | na | na |
| | | **AP51** | na | na | 18,9 | 19,8 | na | na | na | na | na |
| | | **AP52** | na | na | 19,8 | 19,5 | na | na | na | na | na |
| | | **AP53** | na | na | 19,9 | 20,4 | na | na | na | na | na |
| | | **AP54** | na | na | 19,1 | 20,5 | na | na | na | na | na |
| | | **AP55** | na | na | 19,2 | na | na | na | na | na | na |
| | | **AP56** | na | na | 19,0 | na | na | na | na | na | na |
| | | **AP57** | na | na | 19,6 | na | na | na | na | na | na |
| | | **AP58** | na | na | 18,2 | 20,3 | na | na | na | na | na |
| | | **AP59** | na | na | 20,3 | na | na | na | na | na | na |
| | **Skua** (*Stercorarius* sp.) | **SK28** | na | na | 21,5 | 22,7 | na | na | na | na | na |

* FS03 was alive and environmental oro-nasal and peri-anal secretions were sampled in two consecutive days.

### These values refer to environmental oro-nasal secretion swab (indicated as oropharyngeal swab) and peri-anal secretion swab (indicated as cloacal swab).

na: not applicable (not sampled)

nd: not detected

**Supplementary table 3. Correlation of findings for avian cholera between different test types and per tissue.** All animals for which tissues for histopathology were sampled and a negative control Adélie penguin (AP03) are included.

| Organ | Test type | Heroina Island | | | | | | Beagle Island | | | Results (positive/tested) | Negative control |
|---|---|---|---|---|---|---|---|---|---|---|---|---|
| | | Adélie penguin | | | | | Snowy sheathbill | Adélie penguin | | | | |
| | | **AP27** | **AP29** | **AP33** | **AP37** | **AP38** | **SB02** | **AP48** | **AP50** | **AP51** | | **AP03** |
| **Liver** | ***kmt1* qPCR** | na | na | na | ++ | +++ | na | +++ | na | na | **3/3** | - |
| | **H&E** | ++* | na | ++* | - | ++* | na | +++* | +++* | + | **6/7** | - |
| | **Pm ISH** | na | na | na | na | +++ | na | na | +++ | na | **2/2** | - |
| **Brain** | ***kmt1* qPCR** | +++ | +++ | ++ | + | +++ | + | +++ | +++ | +++ | **9/9** | na |
| | **H&E** | - | na | na † | - | ++ | - | ++ | na † | na † | **2/4** | - |
| | **Pm ISH** | na | na | na | na | ++ | - | na | ++ | na | **2/3** | - |
| **Lung** | ***kmt1* qPCR** | +++ | +++ | ++ | + | +++ | ++ | +++ | +++ | +++ | **9/9** | - |
| | **H&E** | - | ++ | - | - | ++ | - | +++ | +++ | na † | **4/8** | - |
| | **Pm ISH** | na | ++ | na | na | ++ | - | na | +++ | na | **3/4** | - |
| **Trachea** | **H&E** | + | + | na | - | - | na | ++ | ++ | na † | **4/6** | - |
| | **Pm ISH** | na | na | na | na | + | na | na | +++ | na | **2/2** | na |
| **Heart** | **H&E** | - | na | + | - | ++ | na | + | ++ | + | **5/6** | - |
| | **Pm ISH** | na | na | na | na | +++ | na | na | na | na | **1/1** | - |
| **Kidney** | **H&E** | - | na | na † | - | +++ | - | na † | ++ | na † | **2/5** | - |
| | **Pm ISH** | na | na | na | na | +++ | na | na | + to +++ | na | **2/3** | - |
| **Spleen** | **H&E** | na | na | na | - | + | na | na | + | na | **2/3** | - |
| | **Pm ISH** | na | na | na | na | + | na | na | +++ | na | **2/2** | - |
| **Intestine** | **H&E** | na | na | na | - | na | + ‡ | - | na | na | **0/3** | na |
| **Evidence of *P. multocida* dissemination beyond liver** | | 3/5 | 3/3 | 3/3 | 2/7 | 6/6 | 2/4 | 4/5 | 6/6 | 3/3 | | 0/7 |

*kmt1* qPCR: DNA loads (40 - Cq): -, not detected; +, 1 to 8; ++, 9 to 16; +++, 17 to 24.
H&E: typical *P. multocida* colonies. -, not observed; +, few observed; ++, observed every 3 high-power fields; +++, observed every high-power field.
Pm ISH: -, not observed; +, few observed; ++, observed every 3 high-power fields; +++, observed every high-power field.
na: not available for analyses
* Microabcesses present
† Tissues sampled, but excluded from analyses because of advanced post mortem changes.
‡ The digesta in the intestinal lumen tested positive for *P. multocida* ISH, but intestinal tissue did not.

**Supplementary Table 4. Metadata, sequencing, MLST and serotyping data of the *Pasteurella multocida* isolates from animals found dead from Hope Bay, Heroina Island, Beagle Island and Paulet Island.** All the locations where positive animals for *Pasteurella multocida* were found (6) are included here. Only samples from which a chromosome completeness ≥60% could be obtained are included.

| Animal ID | Site of collection | Species | Age class | Date of collection (dd-mm-yy) | Sample type | *Pasteurella multocida* reads (Kraken2) | Chromosome completeness | MLST | | | | Capsular type | LPS group | GenBank accession |
|---|---|---|---|---|---|---|---|---|---|---|---|---|---|---|
| | | | | | | | | RIRDC | | Multi-host | | | | |
| | | | | | | | | ST | Nearest ST* | ST | Nearest ST* | | | |
| **BSK01** | Hope Bay | *Stercorarius antarcticus* | Adult | 18-03-24 | Liver | 37900 | 93.4% | 61 | - | 91 | - | A | L1 | *ongoing* |
| **SK06** | Hope Bay | *Stercorarius* sp. | Adult | 18-03-24 | Liver | 145000 | 94.9% | 61 | - | 91 | - | A | L1 | *ongoing* |
| **SK18** | Heroina Is. | *Stercorarius* sp. | Adult | 25-03-24 | Lung | 18000 | 67.7% | nd | 51, **61**, 158 | nd | **91**, 94, 195 | nd | L1 | *ongoing* |
| **BSK20** | Heroina Is. | *Stercorarius antarcticus* | Adult | 25-03-24 | Lung | 11700 | 69.7% | nd | 60, 279, **61**, 507, 452, 278, 158, 51 | nd | **91**, 94, 195 | nd | L1 | *ongoing* |
| **SB03** | Heroina Is. | *Chionis albus* | Adult | 25-03-24 | Brain | 31200 | 91.7% | nd | 51 | nd | **91**, 121 | A | L1 | *ongoing* |
| **AP28** | Heroina Is. | *Pygoscelis adeliae* | Adult | 25-03-24 | Brain | 8820 | 62.4% | nd | 61 | nd | 91 | nd | L1 | *ongoing* |
| **AP38** | Heroina Is. | *Pygoscelis adeliae* | Adult | 25-03-24 | Liver | 29400 | 92.5% | 61 | - | nd | **91**, 121 | A | L1 | *ongoing* |
| **AP39** | Heroina Is. | *Pygoscelis adeliae* | Adult | 25-03-24 | Lung | 17100 | 85.3% | 61 | - | nd | 91 | nd | L1 | *ongoing* |
| **AP50** | Beagle Is. | *Pygoscelis adeliae* | Adult | 26-03-24 | Lung | 10000 | 65.1% | nd | 61 | nd | 91 | A | L1 | *ongoing* |
| **AP51** | Beagle Is. | *Pygoscelis adeliae* | Adult | 26-03-24 | Lung | 11800 | 74.1% | nd | 61 | nd | 91 | A | L1 | *ongoing* |
| **AP52** | Beagle Is. | *Pygoscelis adeliae* | Adult | 26-03-24 | Brain | 12900 | 77.7% | 61 | - | nd | 91 | nd | L1 | *ongoing* |
| **AP57** | Beagle Is. | *Pygoscelis adeliae* | Adult | 26-03-24 | Brain | 17900 | 84.8% | nd | 61 | nd | 91 | nd | L1 | *ongoing* |
| **AP58** | Beagle Is. | *Pygoscelis adeliae* | Adult | 26-03-24 | Lung | 12900 | 76.9% | nd | 61 | nd | 91 | A | L1 | *ongoing* |
| **SK29** | Paulet Is. | *Stercorarius* sp. | Adult | 27-03-24 | Lung | 17200 | 82.7% | 61 | - | nd | **91**, 121 | A | L1 | *ongoing* |

* It indicates the closest ST assigned by the server based on the quality of the query sequences. The server could not assign a precise ST due to imperfect match with the reference sequence or low sequence coverage due to limited chromosome completeness.

nd: not determined.

**Supplementary table 5. Levels of confidence of diagnoses.** Categories were applied to each diagnosis to indicate the confidence level, taking into consideration the lack of disease history for the sampled animals. As GAPDH or β-actin RNA was detected in all the samples, all the real-time PCR results could be considered regardless of the results of these RT-qPCRs.

| Category | Samples available for testing | Level of matching of results |
|---|---|---|
| **High** | Sample available both for (RT-)qPCRs and microscopy | (RT-)qPCR result(s) matches microscopy result |
| **Medium** | Sample available both for (RT-)qPCRs and microscopy | (RT-)qPCR result(s) does not match microscopy result |
| | Sample available for (RT-)qPCRs but not for microscopy | Matching of (RT-)qPCR result with microscopy result is not possible |
| **Low** | Sample available for (RT-)qPCRs but not for microscopy, with low Cq values or not detected by (RT-)qPCR | Matching of (RT-)qPCR result with microscopy result is not possible |
| **Inconclusive** | (RT-)qPCRs negative or close to the limit of detection, samples for microscopy not available, or no lesions detected that can explain disease leading to death. | As there are no positive results, matching is not applicable |
